# Optimizing AAV2/6 microglial targeting identified enhanced efficiency in the photoreceptor degenerative environment

**DOI:** 10.1101/2021.06.02.442814

**Authors:** Margaret E Maes, Gabriele M Wögenstein, Gloria Colombo, Raquel Casado-Polanco, Sandra Siegert

## Abstract

Adeno-associated viruses (AAVs) are widely used to deliver genetic material *in vivo* to distinct cell types such as neurons or glial cells allowing for targeted manipulation. Transduction of microglia is mostly excluded from this strategy likely due to the cells’ heterogeneous state upon environmental changes, which makes AAV design challenging. Here, we established the retina as a model system for microglial AAV validation and optimization. First, we show that AAV2/6 transduced microglia in both synaptic layers, where layer preference corresponds to the intravitreal or subretinal delivery method. Surprisingly, we observed significantly enhanced microglial transduction during photoreceptor degeneration. Thus, we modified the AAV6 capsid to reduce heparin binding resulting in increased microglial transduction in the outer plexiform layer. Finally, to improve microglial-specific transduction, we validated a Cre-dependent transgene delivery cassette.Together, our results provide a foundation for future studies optimizing AAV-mediated microglia transduction and highlight that environmental conditions influence microglial transduction efficiency.

## Introduction

Viral vector engineering has become an effective strategy for *in vivo* delivery of genetic material to distinct cell populations. Due to their ease of engineering and production ^1^, adeno-associated viruses (AAV) are widely used to target various cell types of the central nervous system (CNS) ^2^. However, microglia, the resident immune cell of the CNS, are excluded from this success. The occasionally reported microglial transduction *in vivo* ^3–5^ is rather inefficient and limits microglial manipulation *in vivo.* Yet, we need strategies to selectively alter microglia to obtain knowledge about their function within local environments, and to identify their impact in disease onset and progression ^6, 7^. So far, the lack of robust and systematic investigation of *in vivo* targeting strategies has hindered viral delivery optimization for microglia and is a knowledge gap that needs to be addressed.

Successful viral transduction strategies depend on maximizing cell-specific targeting and minimizing off-target gene expression. By combining known cellular tropism of AAV capsid serotypes with cell type-selective promoters, this goal has been met for many neurons and glia. For microglia, neither an *in vitro* screen of ten capsids with a constitutive promoter ^8^, nor an *in vivo* screen of three capsids and over 200 synthetic promoters ^8, 9^ led to sufficient microglia transduction ^5^. So far, the most encouraging microglia-targeting AAV involved an engineered AAV2/6^TYY^ capsid, which contains mutations to prevent AAV proteasomal degradation upon entry into the target cell ^10^ combined with the promoter, cluster of differentiation 68 (CD68). This AAV2/6^TYY^ was reported to target hippocampal microglia *in vivo* ^8^, although, the *in vitro* transduction efficiency was much higher. This discrepancy could be explained by the difference between *in vitro* and *in vivo* microglial transcriptional signature ^11–13^. Therefore, optimization of AAV to transduce microglia should be performed in an *in vivo* setting, which requires an anatomically defined and well-controlled environment. The retina provides an ideal model, as its highly ordered structure demarcates two synaptic layers, each occupied by distinct microglial niches ^14, 15^. Viral delivery is fast, minimally invasive, and several well-characterized degenerative disease models are available, along with known disease-associated microglial genes ^6^.

Here, we first assessed the feasibility for *in vivo* AAV-mediated microglial transduction in the retina using scAAV2/6^TYY^-CD68-GFP. We systematically investigated different viral delivery strategies and whether a degenerative environment affected microglia susceptibility to AAV transduction. Surprisingly, microglial transduction improved across the plexiform layers when degeneration was initiated in the outer retinal layer. Based on this finding, we engineered an AAV capsid to promote spread within the outer retina in a non-degenerative environment, and confirmed its selective transduction in microglia of the OPL. Finally, we optimized the AAV specificity to reduce off-target expression in other retinal cells by combining a double-inverted transgene cassette with a Cre-mouse line. Overall, this work established the retina as a model system to validate future AAV modification for microglial transduction, which will be relevant for application in other brain regions.

## Results

### Viral delivery route corresponds with layer-specific microglial transduction and viral spread

To investigate whether retinal microglia can be successfully transduced with AAV, we took advantage of scAAV2/6 ^TYY^-CD68-GFP, which consists of a modified AAV6 ^TYY^ capsid and a transgene encoding for GFP driven under the monocyte and tissue macrophage-selective CD68 promoter (**Supplementary Fig. 1a**) ^8^. CD68 transcripts are reliably found *in vivo* in brain microglia ^16^, as well as in the adult retina ^6^ (**Supplementary Fig. 1b**). Furthermore, a 5’ mutated inverted terminal repeat (ITR) flanks the AAV2 genome for self-complementary (sc) assembly and faster transgene expression ^8^. After scAAV2/6 ^TYY^-CD68-GFP production, we confirmed GFP expression in microglia of primary mixed glial culture *in vitro* (**Supplementary Fig. 1c**).

Classically in rodent studies, the location of the retinal cell type to be targeted dictates which injection strategy will be used, where subretinal or intravitreal injection preferentially transduces cells in the outer or inner retinal layers, respectively (**Fig. 1a**) ^17–19^. Microglia in the adult retina are localized in both the outer and inner plexiform layer (OPL and IPL, respectively, **Supplementary Fig. 1d**). Thus, to determine which injection method robustly targets OPL_microglia_ and/or IPL_microglia_, we subretinally or intravitreally injected scAAV2/6 ^TYY^-CD68-GFP into adult C57BL6/J mice (**Fig. 1a**). After two weeks, we performed immunostaining for the transgene GFP and ionized calcium binding adaptor molecule (Iba1) to label the microglial population ^20^. To assess microglial transduction efficiency, we calculated the ratio of GFP^+^/Iba1^+^cells to the total number of Iba1^+^cells within the region-of-interest 1 (ROI1, **Fig. 1a**). The median transduction efficiency was 6.6% for subretinal and 13% for intravitreal injection with GFP frequently expressed in non-microglia cells (**Supplementary Fig. 1e**). When we quantified the microglial transduction efficiency within OPL or IPL, subretinal injection resulted in higher efficiency of OPL_microglia_ compared to IPL_microglia_ (**Fig. 1b-c**), and *vice versa* for intravitreal injection (**Fig. 1d-e**). To estimate the viral spread throughout the retina, we analyzed the opposing quadrant from the injection site, region-of-interest 2 (ROI2, **Fig. 1a**). Independent from the injection method, the transduction efficiency was significantly reduced between ROI1 and ROI2 (**Fig. 1f**). However, intravitreal injection maintained a slightly higher transduction level for ROI2 suggesting enhanced viral spread.

**Figure 1.**
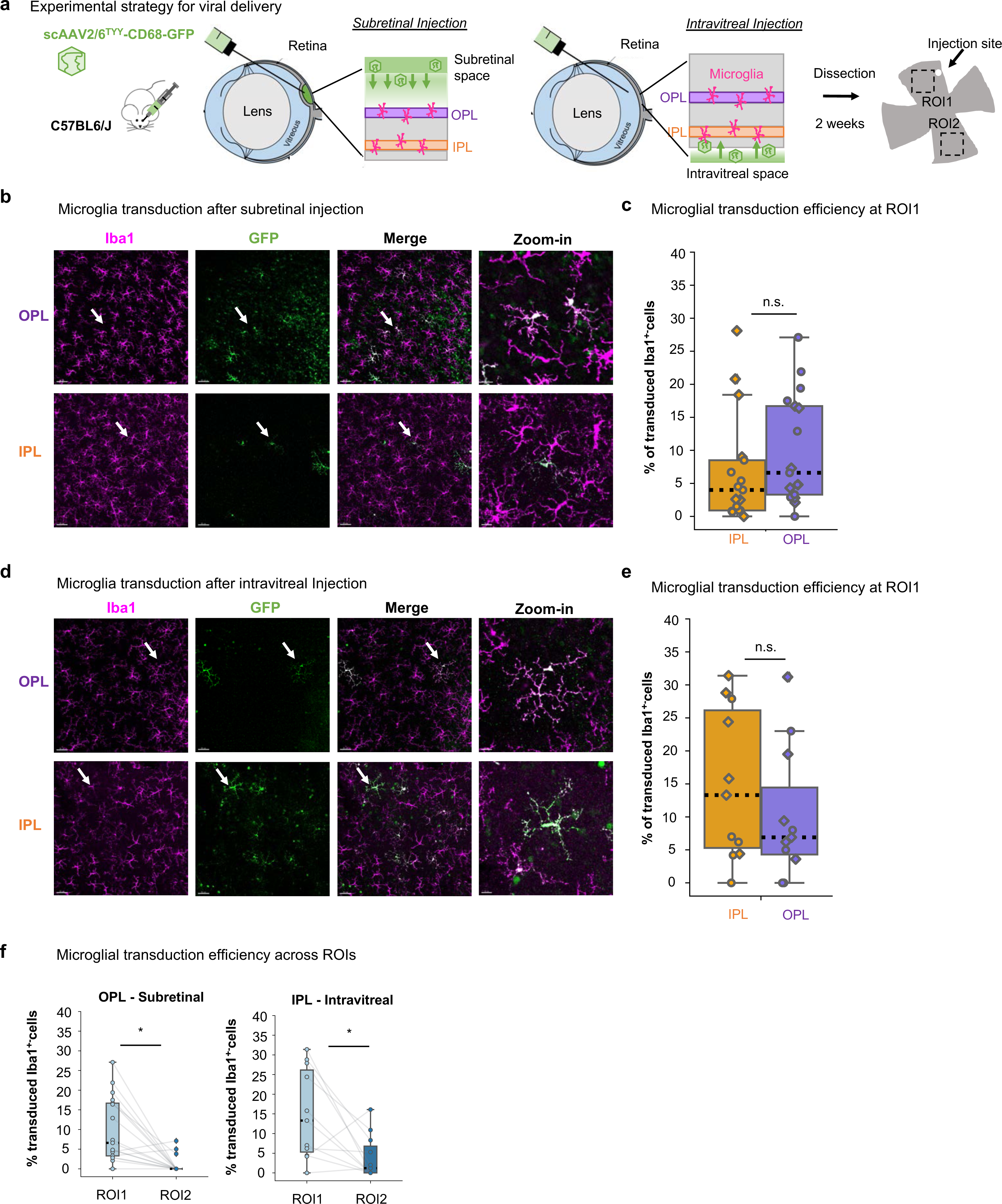
Viral delivery route influences preferred microglial layer transduction. (**a**) Experimental strategy. Adult C57BL6/J mice injected with scAAV2/6^TYY^-CD68-GFP (1*10^12^gc/mL) through subretinal or intravitreal delivery route and collected two weeks later. Images acquired close to the injection site (ROI1), and opposing quadrant (ROI2). (**b-c**) Subretinal, (**d**-**e**) intravitreal injection. **(b**, **d**) Retinal wholemount images of OPL and IPL after subretinal (**b**) and intravitreal (**d**) injection stained with Iba1 (magenta) and GFP (green). White arrows indicate zoom-in region. Scale bar: 50µm, zoom-in: 15µm. (**c**, **e**) Percent microglial transduction efficiency for OPL and IPL microglia at ROI1 after subretinal (**c,** Wilcoxon signed-rank test: *P* = 0.246) and intravitreal (**e,** Wilcoxon signed-rank test: *P* = 0.286) injection. Each point represents ROI1 from one retina. Diamond: male, circle: female. (**f**) Comparison of transduction efficiency across ROIs for individual retinas analyzed in OPL after subretinal (Wilcoxon signed-rank test: *P* = 0.001) or IPL after intravitreal injection (Wilcoxon signed-rank test: *P* = 0.021). Gray lines connect ROIs from a single retina. Subretinal: 17 retinas, 9 mice. Intravitreal: 11 retinas, 6 mice. \**P* < 0.05, ^ns^*P* > 0.05.

Since both viral delivery strategies caused minor injury, thereby initiating microglial proliferation ^21, 22^, we compared the microglial density within ROI1 to naïve, non-injected animals (**Supplementary Fig. 2a**). Only the subretinal method increased microglial density in both plexiform layers. When we assessed each ROI separately, the effect only occurred in ROI1, whereas ROI2 remained at the naïve level (**Supplementary Fig. 2b**). Intravitreal injection did not affect microglial cell density (**Supplementary Fig. 2c**). To confirm that increased efficiency at ROI1 is not due to increased cell density, we calculated the Pearson’s coefficient for both subretinal and intravitreal injections at ROI1 (**Supplementary Fig. 2d**-**e**). We found a negative correlation for subretinal injection, suggesting we may underestimate efficiency, while there was no effect for intravitreal injection.

Together, our data shows that scAAV2/6 ^TYY^-CD68-GFP successfully transduced retinal microglia, preferentially at the ROI closest to the injection site, and that the viral delivery route influences layer preference and viral diffusion across the microglial population.

### Loss of inner retinal barriers did not improve microglial transduction after intravitreal delivery

Effective viral-mediated transgene delivery faces multiple challenges *in vivo*, like physical barriers ^23^. Intravitreally delivered viral particles must first bypass the inner limiting membrane, the dense extracellular matrix around the nerve fiber, and the ganglion cell layer to reach the IPL ^24^. Disrupting these barriers using the optic nerve crush (ONC) model, resulted in greater penetration into the retinal layers for AAV2 ^25–27^. To determine whether ONC influenced microglial transduction, we performed ONC on adult C57BL6/J mice and intravitreally injected scAAV2/6 ^TYY^-CD68-GFP four weeks post-injury (**Fig. 2a**). At this time, microglia have cleared the apoptotic ganglion cells ^28^ and returned to a non-reactive morphology as confirmed by Sholl analysis (**Supplementary Fig. 3a**). Two weeks after the injection, we analyzed the retinas for GFP^+^-microglia and compared to non-crushed, intravitreally injected retinas. Unexpectedly, neither the OPL_microglia_ nor the IPL_microglia_ transduction efficiency improved (**Fig. 2b-e**), even though ONC resulted in a 50% cell loss in the ganglion cell layer (**Supplementary Fig. 3b**). The viral spread was also unchanged in the IPL (**Fig. 2f**). Thus, the ONC-mediated reduction of the physical barrier did not further improve microglial transduction.

**Figure 2.**
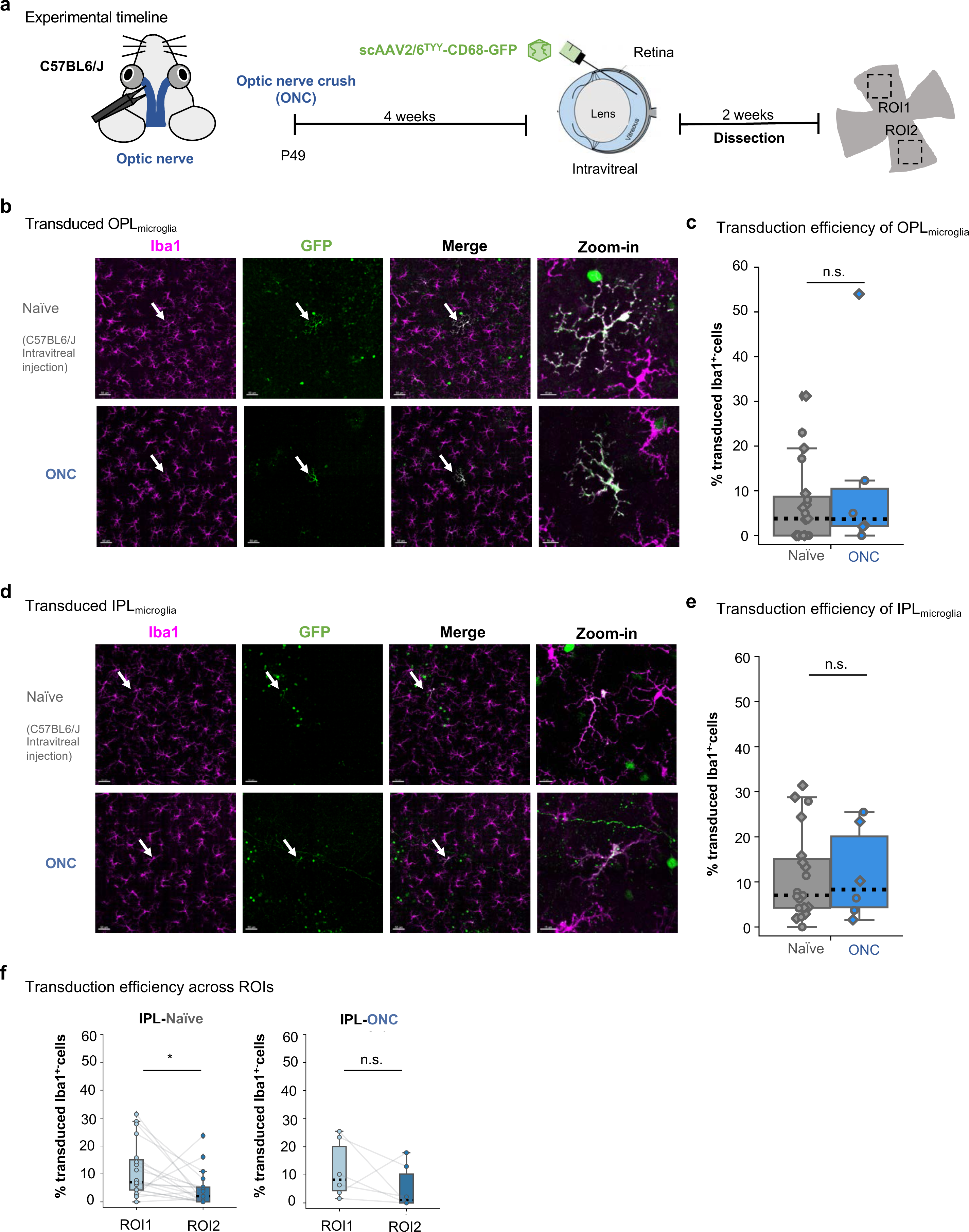
Microglial transduction efficiency was unaltered after optic nerve crush (ONC) (**a**) Experimental timeline. ONC surgery performed on the left eye of adult C57BL6/J mice. scAAV2/6^TYY^-CD68-GFP intravitreally delivered 4 weeks later. Retinas collected 2 weeks after injection. (**b**, **d**) Retinal wholemount images of OPL_microglia_ or IPL_microglia_ after ONC or naïve conditions stained with Iba1 (magenta) and GFP (green). White arrows indicate zoom-in region. Scale bar: 50µm, zoom-in: 15µm. (**c, e**) Percent microglial transduction efficiency for OPL_microglia_ (**c,** Wilcoxon rank-sum test: *P* = 0.633) and IPL_microglia_ (**e,** Wilcoxon rank-sum test: *P* = 0.899) naïve or ONC at ROI1. Each point represents ROI1 from one retina. Diamond: male, circle: female. Two experiments pooled (1*10^12^gc/mL or 1.37*10^11^gc/mL) (**f**) Comparison of transduction efficiency across ROIs for individual retinas analyzed in IPL_microglia_ in naïve (Wilcoxon signed-rank test: *P* = 0.014) or ONC condition (Wilcoxon signed-rank test: *P* = 0.249). Gray lines connect ROIs from a single retina. Naive: n=19 retinas, 12 mice. ONC: n = 6 retinas, 6 mice. \**P* < 0.05, ^ns^*P* > 0.05.

### Rod photoreceptor loss enhanced microglial transduction after subretinal delivery

Subretinally delivered viral particles must pass the densely packed outer nuclear layer (ONL) to target cells in the OPL. Photoreceptor degeneration models such as *rd10*, reduce this physical barrier. *Rd10* harbors a missense point mutation in the *Pde6b* gene leading to progressive rod photoreceptor degeneration ^29^, which peaks at postnatal day 25-30 ^30^. At postnatal day 65 (P65), the ONL thickness was reduced (**Supplementary Fig. 4a**), and microglial density was comparable to naïve (**Supplementary Fig. 4b-c**). Therefore, we subretinally injected scAAV2/6 ^TYY^-CD68-GFP in Pde6b*^rd10/rd10^* mice at P65 and age-matched C57BL6/J controls, and analyzed the retina two weeks post-injection (**Fig. 3a**). We found enhanced OPL_microglia_ transduction in Pde6b*^rd10/rd10^* compared to controls (**Fig. 3b-c**).

**Figure 3.**
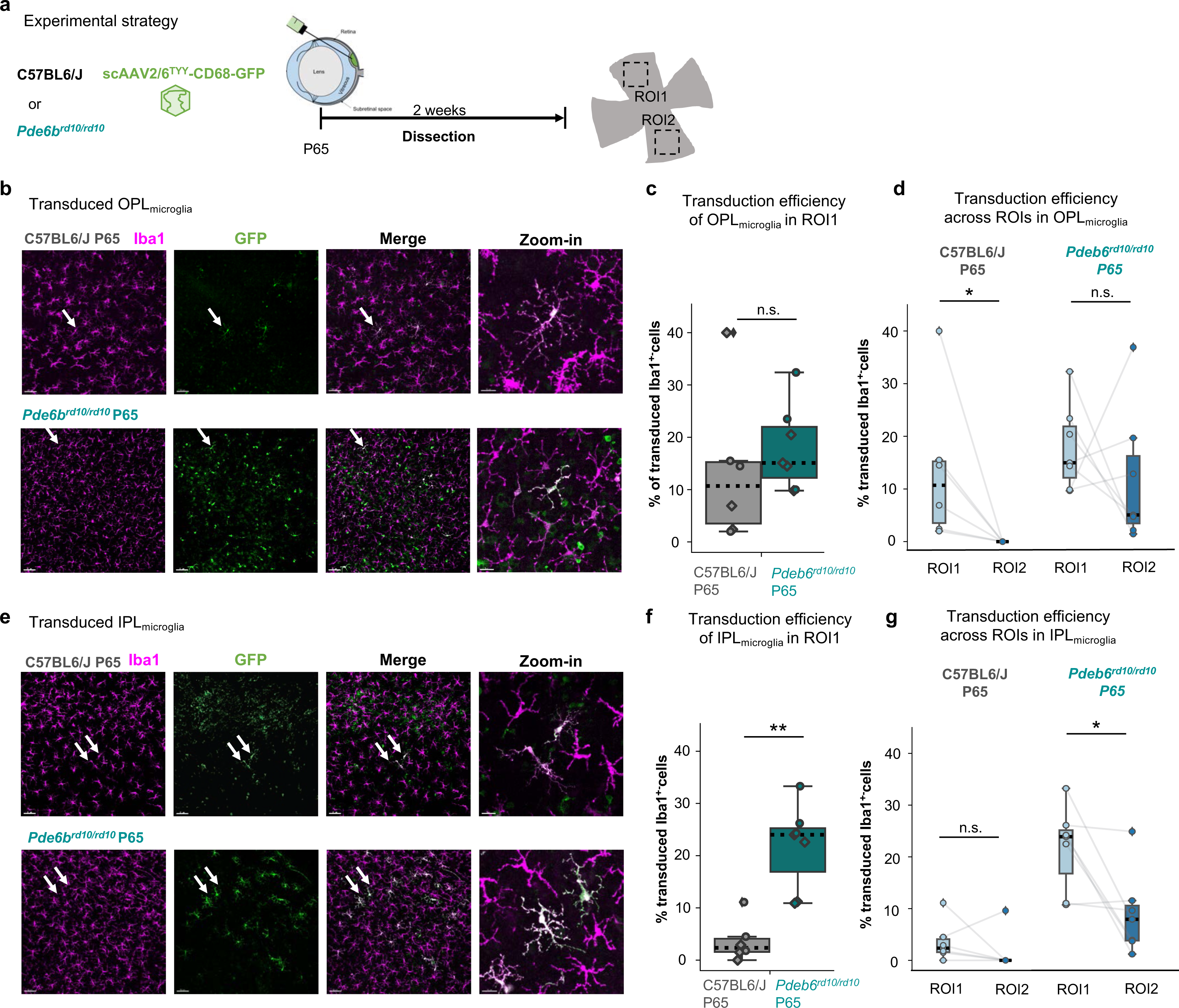
Photoreceptor degeneration increases microglial transduction efficiency and spread throughout retina. (**a**) Experimental timeline. Subretinal delivery of scAAV2/6^TYY^-CD68-GFP (1.37*10^11^gc/mL) to postnatal day 65 (P65) *Pde6b^rd10/rd10^* or C57BL6/J mice. (**b**, **e**) Retinal wholemounts of transduced OPL_microglia_ (**b**) and IPL_microglia_ (**e**) in P65 *Pde6b^rd10/rd10^* retinas immunostained with Iba1 (magenta) or GFP (green). White arrows indicate zoom-in. Scale bar: 50µm, zoom-in: 15µm. (**c**) Comparison of P65 *Pde6b^rd10/rd10^* and C57Bl6/J transduction efficiency OPL_microglia_ (Wilcoxon ranked-sum test, *P* = 0.284) and (**d**) transduction across OPL_microglia_ ROIs (Wilcoxon signed-rank test: C57BL6/J*, P* = 0.027; *Pde6b^rd10/rd10^, P* = 0.398). (**f**) IPL_microglia_, transduction efficiency in P65 *Pde6b^rd10/rd10^* compared to control (Wilcoxon ranked-sum test: *P* = 0.004). (**g**) Transduction across ROIs in IPL_microglia_ (Wilcoxon signed-rank test: C57BL6/J, *P* = 0.345; *Pde6b^rd10/rd10^, P* = 0.027). P65 *Pde6b^rd10/rd10^*: 7 retinas, 4 mice. C57BL6/J: 6 retinas, 4 mice. \*\**P* < 0.005, \**P* < 0.05, ^ns^*P* > 0.05.

Also, the median viral spread significantly improved across ROIs by 5-fold (**Fig. 3d**). Although subretinal delivery is not the optimal route for targeting IPL_microglia_ (**Fig. 1e**), we unexpectedly found a significant increase at both ROIs (**Fig. 3e-g**).

Since ONL loss is apparent by P27 in Pde6b*^rd10/rd10^* (**Supplementary Fig. 4a**), we investigated whether enhanced transduction is already evident. Therefore, we subretinally injected scAAV2/6^TYY-^CD68-GFP (**Supplementary Fig. 5a**). For OPL_microglia_, the transduction efficiency increased, however only for ROI1 (**Supplementary Fig. 5b-d**). For IPL_microglia_, we no longer observed increased transduction efficiency (**Supplementary Fig. 5e-f**), and there was no change in spread across ROIs for either plexiform layer (**Supplementary Fig. 5g**), which is in contrast to Pde6b*^rd10/rd10^* P65.

Taken together, the loss of the physical ONL barrier in Pde6b*^rd10/rd10^* benefits OPL_microglia_ transduction efficiency, while the P65 environment further supports transduction across plexiform layers and ROIs.

### Mutation of AAV6^TYY^ capsid heparin binding sites improved OPL_microglia_ transduction

Extracellular matrix remodeling in the outer retinal layers could explain the increased transduction and spread in the Pde6b*^rd10/rd10^* P65 environment. Boye *et al.* have shown improved outer retinal transduction through selective capsid mutations at binding sites of heparin, a highly-sulfated form of the extracellular matrix component heparan sulfate ^31^. A previously identified single mutation, K531E, reduced heparin binding capacity by AAV6 ^32^. Therefore, we introduced this mutation to the AAV6^TYY^ capsid (scAAV^K531E^-CD68-GFP) and performed subretinal injection in adult C57BL6/J animals. When we analyzed the retinas two weeks later, microglial transduction efficiency did not improve in either plexiform layer (**Supplementary Fig. 6a**). Combined heparin-binding mutations increased viral spread ^31^, thus, we included three additional mutations (R576Q, K493S and K459S, **Fig. 4a**) ^33^. After confirming that the mutated AAV6 capsid (AAV6^Δ4^) transduced microglia in primary mixed glial cells *in vitro* (**Supplementary Fig. 6b**), we subretinally injected scAAV2/6^Δ4^-CD68-GFP into adult C57BL6/J mice (**Fig. 4b**). OPL_microglia_ showed a two-fold increase in transduction for ROI1 compared to scAAV2/6 ^TYY^ (**Fig. 4c-d**). The efficiency did not improve for OPL_microglia_ in ROI2 (**Fig. 4e**), or for IPL_microglia_ in either ROI (**Fig. 4f-h**), suggesting that the AAV2/6^Δ4^ capsid may have limitations in crossing the inner nuclear layer to reach the IPL_microglia_ niche.

**Figure 4.**
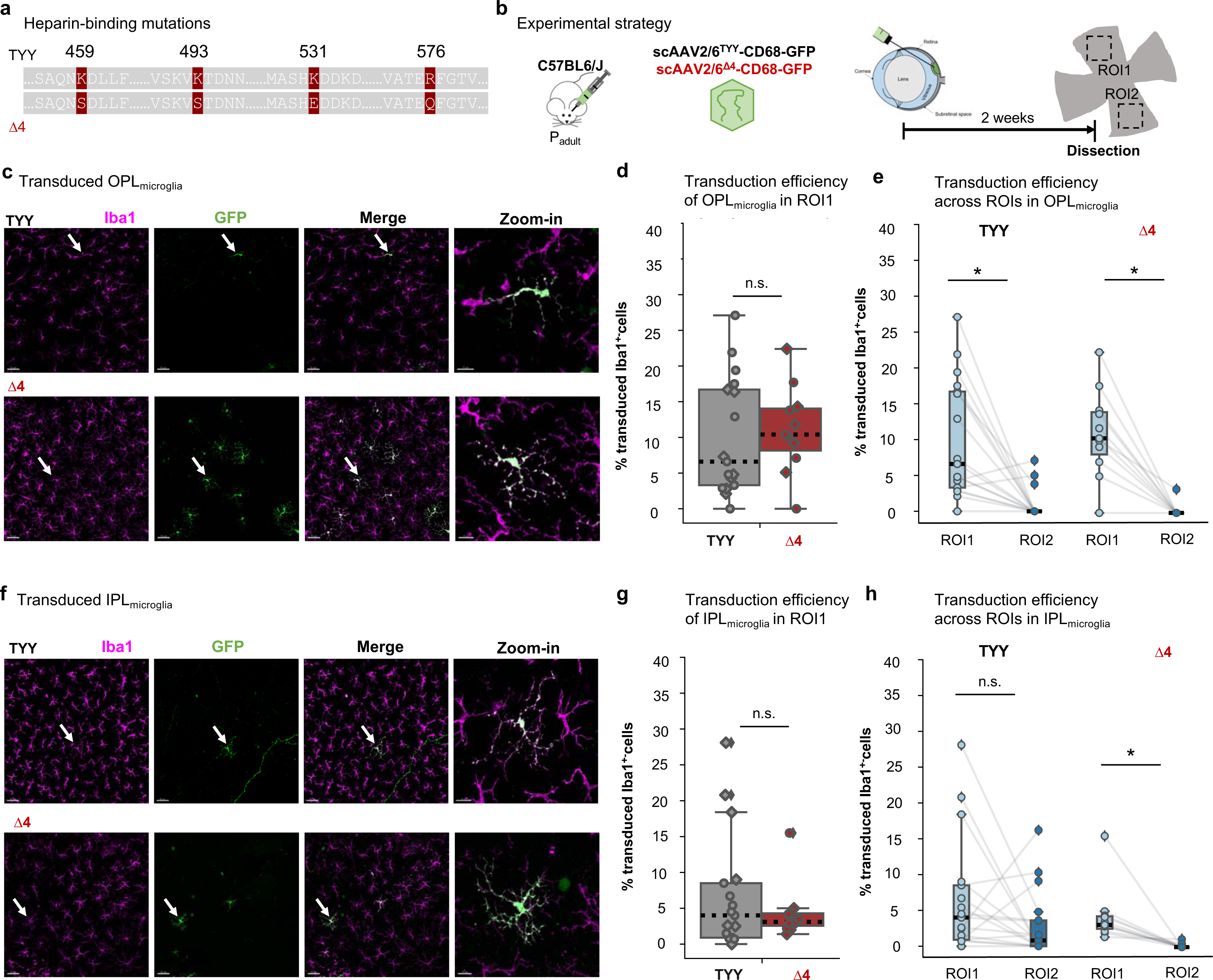
Heparin-binding capsid mutations increase OPL_microglia_ transduction. (**a**) Depiction of the site-specific mutations in the AAV6 capsid (K459S, K493S, K531E, R576Q). (**b**) Experimental timeline. Subretinal delivery of scAAV2/6^TYY^-CD68-GFP or scAAV2/6^Δ4^-CD68-GFP (1*10^12^ gc/mL) to adult C57BL6/J mice. Dissection of retinas followed two weeks later. (**c, f**) Retinal wholemounts immunostained with Iba1 (magenta) and GFP (green) showing transduced OPL_microglia_ and IPL_microglia_ (**f**) in C57BL6/J retinas (dataset from subretinal Fig. 1) with indicated capsid variant. White arrows indicate zoom-in. Scale bar: 50µm, zoom-in: 15µm. (**d**) Comparison of the transduction efficiency in the OPL_microglia_ (Wilcoxon ranked-sum test, *P* = 0.452) and (**g**) IPL_microglia_ (Two sample t-test, *P* = 0.354). (**e**, **h**) Comparison of microglial transduction between ROIs for OPL_microglia_ (**e**, Wilcoxon signed-rank test: TYY, *P* = 0.001; Δ4*, P* = 0.005) or IPL_microglia_ niche (**h**, Wilcoxon signed-rank test: TYY, *P* = 0.092; Δ4*, P* = 0.003). TYY: 17 retinas, 9 mice. Δ4: 11 retinas, 6 mice. \**P* < 0.05, ^ns^*P* > 0.05.

Heparin-binding is required for AAV to pass the inner limiting membrane ^34^, therefore to validate our mutant capsid scAAV2/6^Δ4^-CD68-GFP, we also performed intravitreal injections. Indeed, the percentage of transduced IPL_microglia_ was significantly reduced (**Supplementary Fig. 6c**). We observed few GFP^+^-microglia close to the ROI1 and none in the ROI2 (**Supplementary Fig. 6d**), suggesting minimal access from the injection procedure. Overall, the mutated heparin binding sites in our scAAV2/6^Δ4^-CD68-GFP improved OPL_microglia_ transduction.

### Microglial-specific transduction with Cre recombinase-dependent AAV2/6^Δ4^

Although we improved efficiency with scAAV2/6^Δ4^-CD68-GFP, we still faced off-target transgene expression frequently in cone photoreceptors in the outer nuclear (ONL), Müller glia in the inner nuclear (INL), and ganglion cells and astrocytes in the ganglion cell layer (GCL) (**Supplementary Fig. 7a-c**). When we counted the number of retinas with at least one positive off-target cell, all retinas in INL and GCL exhibited off target cells, and over half in the ONL (**Supplementary Fig. 7d**). Since this unspecific expression limits downstream applications and due to the lack of synthetic microglial-selective promoters ^9^, we decided to improve specificity introducing an inducible double-floxed inverse orientation (DIO) sequence into the transfer vector (**Fig. 5a**). We subretinally injected the scAAV2/6^Δ4^-CD68-DIO-GFP into Cx3cr1^CreERT2/+^ mice, which express tamoxifen-inducible Cre recombinase under the Cx3cr1 promoter ^35^. To induce Cre-mediated inversion, we intraperitoneally injected tamoxifen for three consecutive days one week after viral delivery ^36^, and analyzed transgene expression two weeks later (**Fig. 5b**).

**Figure 5.**
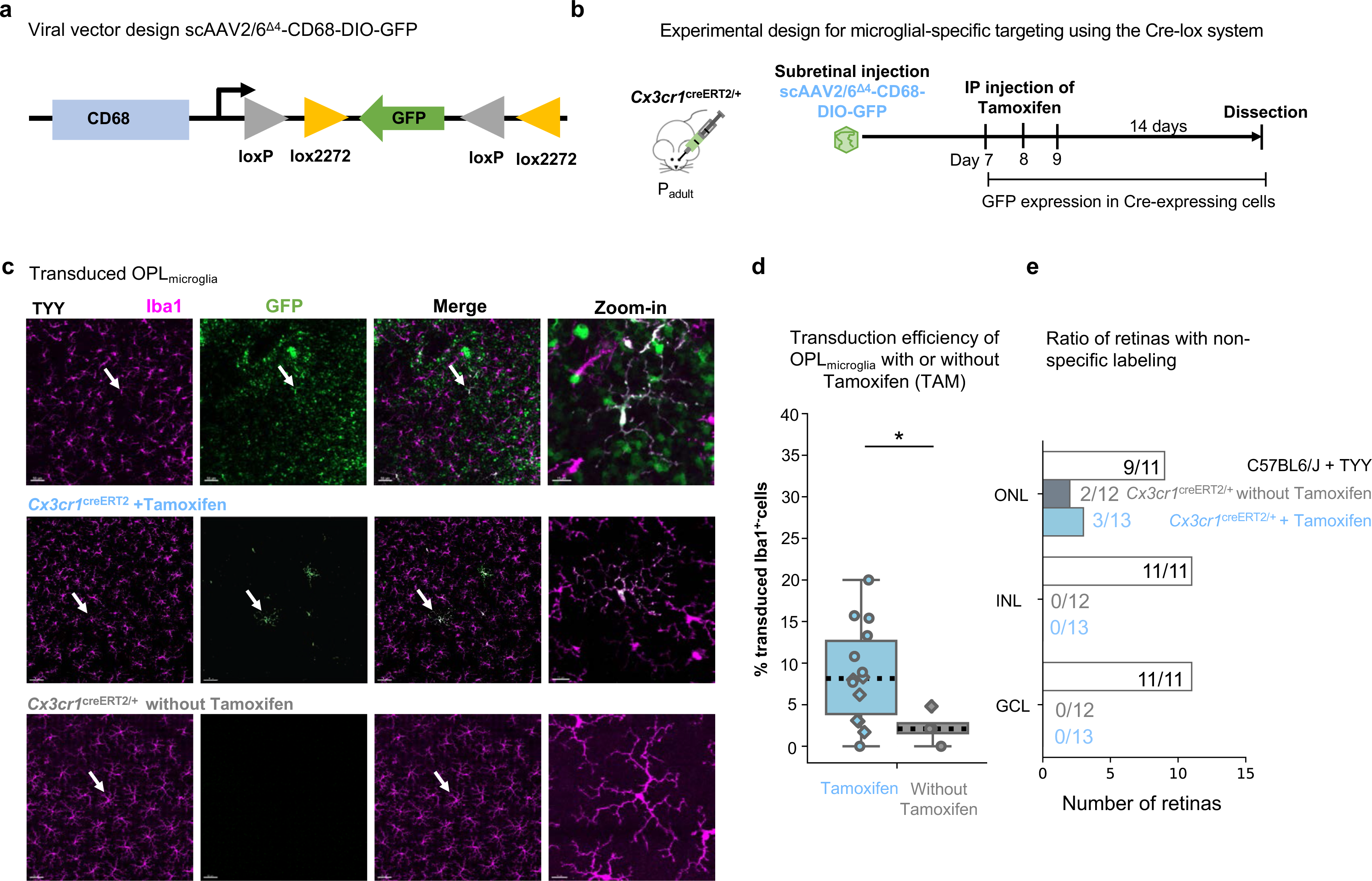
Microglia-specific transgene expression using Cre-dependent scAAV2/6^Δ4^. (**a**) Transfer vector design. Two loxP sites flank the inverted GFP transgene. (**b**) Experimental timeline. Adult Cx3cr1^creERT2/+^ mice received subretinal injection of scAAV^Δ4^-CD68-DIO-GFP (3*10^12^ gc/mL) and tamoxifen injections for three consecutive days, one week after viral injection. Two weeks later retinas were collected. (**c**) Retinal wholemounts of transduced OPL_microglia_ of C57BL6/J mice after subretinal injection of scAAV^TYY^-CD68-DIO-GFP (Fig. 1) and Cx3cr1^creERT2/+^ mice after subretinal injection of scAAV^Δ4^-CD68-DIO-GFP with and without receiving Tamoxifen treatment. White arrows indicate zoom-in. Scale bar: 50µm, zoom-in: 15µm. (**d**) Comparison of transduction efficiency in the OPL with and without Tamoxifen treatment. (Wilcoxon ranked-sum test, *P* =0.038). (**e**) Ratio of analyzed retinas showing off-target GFP expression in the indicated retinal layers. Dataset for comparison. Cx3cr1^creERT2/+^ with Tam: 14 retinas, 9 mice. Cx3cr1^creERT2/+^ without Tam: 4 retinas, 3 mice. \**P* < 0.05, ^ns^*P* > 0.05.

As expected, GFP expression was selective for microglia and tamoxifen-dependent (**Fig. 5c-d**). Without tamoxifen, we occasionally observed microglial transduction (**Fig. 5d**), which is expected based on previously reported ‘Cre-leakiness’ in the Cx3cr1^CreERT2^ model ^37^.

Furthermore, the absence of the Cre recombinase in C57BL6/J animals prevented DIO-mediated inversion and GFP-expression (**Supplementary Fig. 8a**).

Minimal off-target transgene expression remained with the combination of DIO-AAV and Cx3cr1^CreERT2/+^ mice. Within ROI1, only the ONL showed 1-2 off-target cells in 3 out of 13 retinas (**Fig. 5e**). This could be attributed to spontaneous transgene inversion during AAV production ^38^. Indeed, we observed similar off-target expression in the ONL of both Cx3cr1^CreERT2/+^ mice without tamoxifen and C57BL6/J with tamoxifen (**Fig. 5d**).

Together, the combined approach using scAAV2/6^Δ4^-CD68-DIO-GFP and Cx3cr1^CreERT2/+^ mice confirms microglial-specific transgene expression with minimal off-target labeling.

### Optimization of microglial transduction using Cre-dependent AAV2/6^Δ4^

Using the retina as a model environment to establish and validate *in vivo* microglia-specific targeting, it allowed us to further refine technical aspects of our system and explore additional biological questions.

One technical challenge could be cross-recombination when using the Cre-dependent AAV in tandem with a floxed reporter mouse line ^39^. To estimate the likelihood of cross-recombination which can result in loss of the reporter transgene expression, we subretinally injected scAAV2/6^Δ4^-CD68-DIO-GFP into Cx3cr1^CreERT2/+^/Rosa26^Ai9/+^ tdTomato reporter mice ^40^. In this mouse model, microglia express tdTomato upon tamoxifen injection (**Fig 6a**). Less than 5% of the transduced GFP^+^ microglia lacked tdTomato expression in both plexiform layers (**Fig. 6b**), suggesting that cross-recombination occurs very rarely in this reporter line.

**Fig. 6.**
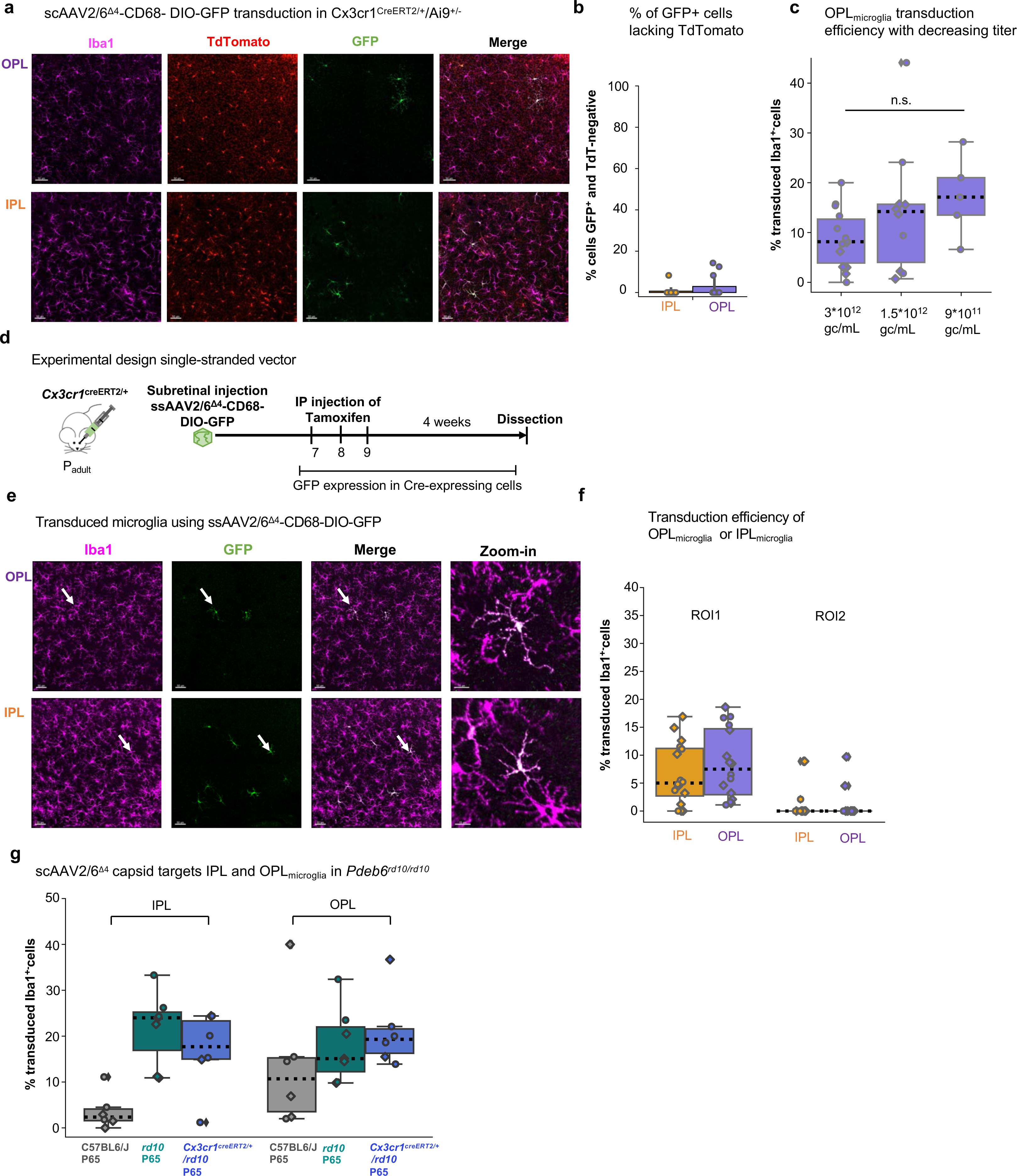
Optimization of microglia-targeting using AAV2/6^Δ4^-CD68-DIO-GFP. (a) Retinal wholemounts of the OPL_microglia_ and IPL_microglia_ of Cx3cr1^CreERT2/+^/Ai9^+/-^ mice after subretinal injection of scAAV^Δ4^-CD68-DIO-GFP and tamoxifen treatment. Scale bar: 50µm. (b)) Quantification of co-expression of GFP and TdTomato. (**c**) Comparison of OPL_microglia_ transduction in using different viral titers. (Kruskal-wallis test: *P* =0.127) (**d**) Experimental timeline. Cx3cr1^creERT2/+^ mice received subretinal injection of ssAAV^Δ4^-CD68-DIO-GFP and Tamoxifen injections for three consecutive days one week after viral injection. 4 weeks after the first Tamoxifen injection the retinas were collected. (**e**) Retinal wholemounts of OPL_microglia_ and IPL_microglia_ of Cx3cr1^creERT2/+^ mice after subretinal injection of ssAAV^Δ4^-CD68-DIO-GFP and Tamoxifen treatment. White arrows indicate zoom-in. Scale bar: 50µm, zoom-in: 15µm. (**f**) Comparison of microglial transduction efficiency between ROIs for both OPL and IPL niche after subretinal delivery of ssAAV2/6^Δ4^-CD68-GFP. (**g**) Comparison of transduction efficiency in the OPL_microglia_ and IPL_microglia_ of C57BL6/J, *Pde6b^rd10/rd10^* (rd10) (Figure 3 dataset) and Cx3cr1^creERT2/+^/*Pdeb6^rd10/10^* mice after subretinal virus delivery. \**P* < 0.05, ^ns^*P* > 0.05.

Another technical consideration is viral titer, which can influence transduction efficiency *in vivo* ^41^. Thus, we compared how subretinal injection of different viral titers impacted OPL_microglia_ transduction efficiency using the scAAV2/6^Δ4^-CD68-DIO-GFP in adult Cx3cr1^CreERT2/+^ animals. A viral titer of 3*10^12^ genome copies/mL resulted in 8% OPL_microglia_ transduction (**Fig. 6c**), while halving the titer to 1.5*10^12^ genome copies/mL, led to 14% microglial transduction. Further reduction of the injection volume to a titer of 9*10^11^ genome copies/mL slightly enhanced OPL_microglia_ transduction to 17%, suggesting that lower titer might be beneficial for subretinal injections.

One major limitation of self-complementary AAVs is the reduced packaging size, which enhances transgene expression ^42^, but on the other hand, halves the AAV packaging size. Thus, we generated a single-stranded AAV (ssAAV) vector and subretinally injected ssAAV2/6^Δ4^-CD68-DIO-GFP into the retina. The overall transduction efficiency was consistent with the scAAV2/6^Δ4^-CD68-DIO-GFP (**Fig. 6d-f**), and can allow a size up to 3 kb for the transgene after exclusion of promoter, ITR and polyA sequences.

Finally, the retina allows for investigation of niche-selective microglial targeting. Our AAV2/6^Δ4^-capsid showed preference for OPL_microglia_, while IPL_microglia_ transduction remained low (**Fig. 4g**, **Supplementary Fig. 8b-c**). Since we found a significant increase in IPL_microglia_ transduction in the P65 Pde6b*^rd10/rd10^* condition, we questioned whether a potential niche-selective expression of the scAAV2/6^Δ4^-capsid remained (**Fig. 3c**). Thus, we crossed the Cx3cr1^CreERT2/+^ onto the Pde6b*^rd10/rd10^* background, subretinally injected scAAV2/6^Δ4^-CD68-DIO-GFP, and compared the results to scAAV2/6 ^TYY^-CD68-GFP from **Fig. 3c**. ScAAV2/6^Δ4^ transduced IPL_microglia_ at a similar level to scAAV2/6^TYY^ (**Fig. 6g**), suggesting that the AAV2/6^Δ4^ capsid is capable of transducing both retinal microglial niches.

Taken together, we demonstrated the efficacy of the Cre-dependent AAV2/6^Δ4^, and validated the viral tool for microglial transduction in the retina.

## Discussion

In this study, we show for the first time that retinal microglia can be successfully targeted using AAV, and is influenced by delivery route. We identified that during photoreceptor degeneration, microglial transduction improved and used this result to inform a modified AAV2/6^Δ4^ that enhanced OPL_microglia_ in adult animals. Finally, we optimized several parameters to improve AAV2/6^Δ4^ for future microglial transduction studies.

### Transduction of microglia in degenerative conditions

In two retinal degenerative conditions, we were able to use known environmental changes to dissect parameters that may affect microglial transduction efficiency. Optic nerve crush (ONC) disrupts the inner limiting membrane and reduces the nerve fiber layer thickness ^25^. We expected that this would improve AAV2/6 access to reach the retinal layers and therefore microglial transduction, however this was not the case. One explanation could be the difference in AAV2 and AAV6 capsid tropism and heparin binding affinity. AAV2 capsid has a high affinity for heparin and accumulates at the inner limiting membrane, which requires HSPG-binding to pass through ^43, 44^. In contrast, AAV6 uses both HSPG and sialic acid for cellular attachment and entry, and displays a weaker heparin binding affinity than AAV2 capsid ^32, 45^. This suggests that AAV6 is inferior to AAV2 at spreading evenly throughout the retina after intravitreal delivery ^46, 47^, and therefore, it may not gain an advantage to targeting cells after ONC.

In contrast to ONC, subretinal injection in the Pde6b*^rd10/rd10^* resulted in a robust microglial transduction (**Fig. 3**). The virus is trapped in the subretinal space, where the weaker heparin binding affinity of AAV2/6 ^32, 45^ may assist its propagation through the subretinal space before cell attachment. Furthermore, extracellular matrix remodeling, accompanied by the reduction of the densely packed ONL, could affect viral cell attachment and transduction in the subretinal space ^48^.

The surprising finding was the significant increase in IPL_microglia_ transduction in the P65 Pde6b*^rd10/rd10^* environment (**Fig. 3**). The reason for this could be two-fold: either the extracellular matrix has already been restructured in the INL, allowing easier access of AAV to the IPL_microglia_, and/or a multifaceted glial response has adapted to the changing degenerative environment. Microglia are known to take on new and distinct transcriptional states in degenerative conditions ^15, 49^. The increase in IPL_microglia_ transduction from P27 to P65 Pde6b*^rd10/rd10^* could suggest that microglia become more susceptible to transduction throughout disease progression, which is an interesting observation for follow-up studies.

### Nonspecific Labeling

Off-target transgene expression is an on-going challenge in viral gene delivery, and can only be circumvented by optimizing viral tropism and cell type-specific promoters. Both aspects are ill-defined for microglia. Besides AAVs, lentiviral vectors have been used to target microglia, but they prevent off-target transgene expression upon employing a microRNA-9 sponge ^50^. This strategy is suboptimal because microRNA-9 has known effects on neurogenesis and synaptic plasticity ^51, 52^, thus sequestering microRNA-9 in off-target cells could result in unknown effects that confound experimental results. Here, we have focused on AAV2/6 due to its suggested specificity in targeting microglia *in vivo* ^8^. However, we found that scAAV2/6^YTT^-CD68-GFP resulted in strong off-target cell type labeling in the retina.

Implementing a tamoxifen-inducible system combining Cx3Cr1^CreERT2^ animals with scAAV2/6^Δ4^-DIO-CD68-GFP led to microglia-specific labeling in the GCL and INL (**Fig. 5i**). Only in the ONL, we found non-specific labeling of 1-5 cells per ROI in only a few retinas analyzed. We suspect that this is due to spontaneous inversion of DIO transgenes ^38^, which we substantiated in our control experiments (**Fig. 5d**). To eradicate off-target expression, incorporation of mutant recombinase recognition sites can prevent the spontaneous inversion ^38^.

Nevertheless, the microglial community is in need of microglia-specific promoters. The rapid environmental adaptation of microglia is reflected by transcriptional changes, which makes it challenging to identify reliable promoters that encompass all potential microglial conditions. Promoters such as CD68 and Cx3cr1 also label blood-derived or brain-barrier-associated macrophages ^53, 54^. Proposed new markers from RNA sequencing studies such as transmembrane protein 119 (TMEM119) have been recently challenged on their specificity *e.g.* in the retina ^55^. On the other hand, the retina provides a unique opportunity to explore novel viral transduction strategies and validating new promoters in future studies ^9^.

### Variability of microglial transduction efficiency

Across CNS regions and viral types, microglial transduction remains at low efficiencies and variable within conditions ^5^. We observed variation within experimental groups throughout our work, which was independent from injection method, capsid variants, Cre-dependent genomes or animal sex. These variations appear in other studies, yet without further discussion of the source of the variation ^56, 57^.

Transcriptomic studies have made it clear that the microglial population is highly heterogeneous ^49, 58^. On the one hand, microglia maintain a common gene signature, which overlaps with other immune cells, yet on the other hand, microglia adapt to their local CNS niche, which might manifest into more distinctly heterogenous populations after damage ^15, 59^. Since we know microglia respond to both the injection-induced insult and to the AAV particles themselves ^22^, we may create a new microglial niches, which contribute to an overall increase in heterogeneity across the microglial population. One way to mitigate these effects could be to deliver the virus alongside compounds known to reduce microglia reactivity, for example, TSPO ^60^, or minocycline ^61^, however the effects on the experimental condition must also be considered.

### Conclusion

Our work highlights the feasibility of microglial transduction in the retina with a modified AAV2/6. Using retinal degeneration models to assess the effect of altered environments on microglial transduction, we found enhanced microglia targeting in a photoreceptor degenerative model. We applied this finding to generate a modified AAV2/6 that increases OPL_microglia_ transduction in healthy adult animals. Finally, we validated and optimized a Cre-dependent AAV strategy for specific microglial targeting *in vivo*, that provides the foundation for future studies.

## Materials and Methods

### Cloning

The self-complementary transfer vector (pAAV2-CD68-MVMi-DIO-eGFP) was generated using the pAAV2-CD68-MVMi-hGFP plasmid kindly provided by Rosario *et. al.* and an RV-CAG-DIO-GFP plasmid, purchased from Addgene (#87662). Both plasmids were digested with *Sac*I and *Pst*I (New England BioLabs) to obtain fragments containing the vector backbone CD68 promoter and the DIO-eGFP insert, which were ligated to yield the final product pAAV2-CD68-MVMi-DIO-eGFP. The single-stranded (ss) pAAV2-CD68-MVMi-DIO-eGFP plasmid was generated by PCR amplification of the insert containing CD68 promoter and loxP sites flanking the eGFP using primers CD68-DIO-eGFP For and CD68-DIO-eGFP Rev (Table 1) from the pAAV2-CD68-MVMi-DIO-eGFP plasmid. The resulting product along with ssAAV backbone (pAAV-ProA3(SynP137)-ChR2d-eGFP-WPRE), kindly provided by Botond Roska, was digested with *Mlu*I and *Rsr*II (New England BioLabs, R3198S, R0501S). The ligated product was transformed and purified, then confirmed by Sanger sequencing.

**Table 1:**
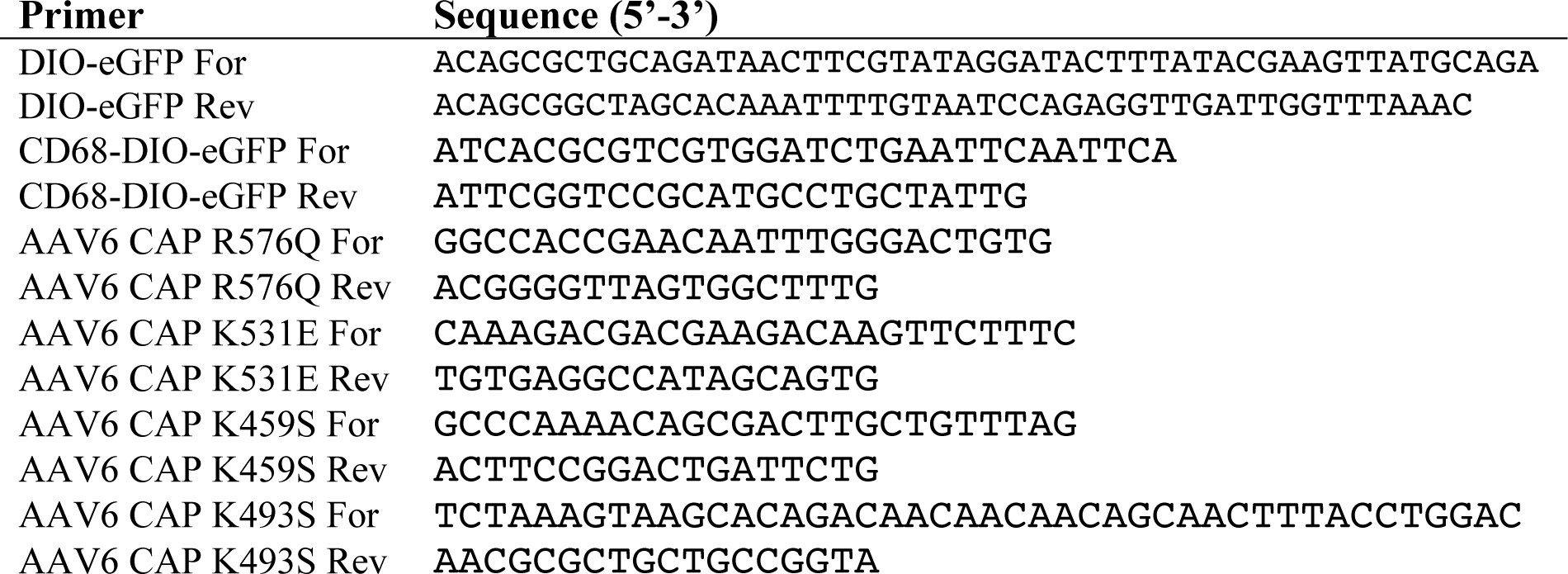
List of primer pairs used for insert amplification to generate the different AAV2 transfer vectors and to perform side directed mutagenesis of the plasmid pACGr2c6-T492V-Y705F-Y731F

**Table 2:**
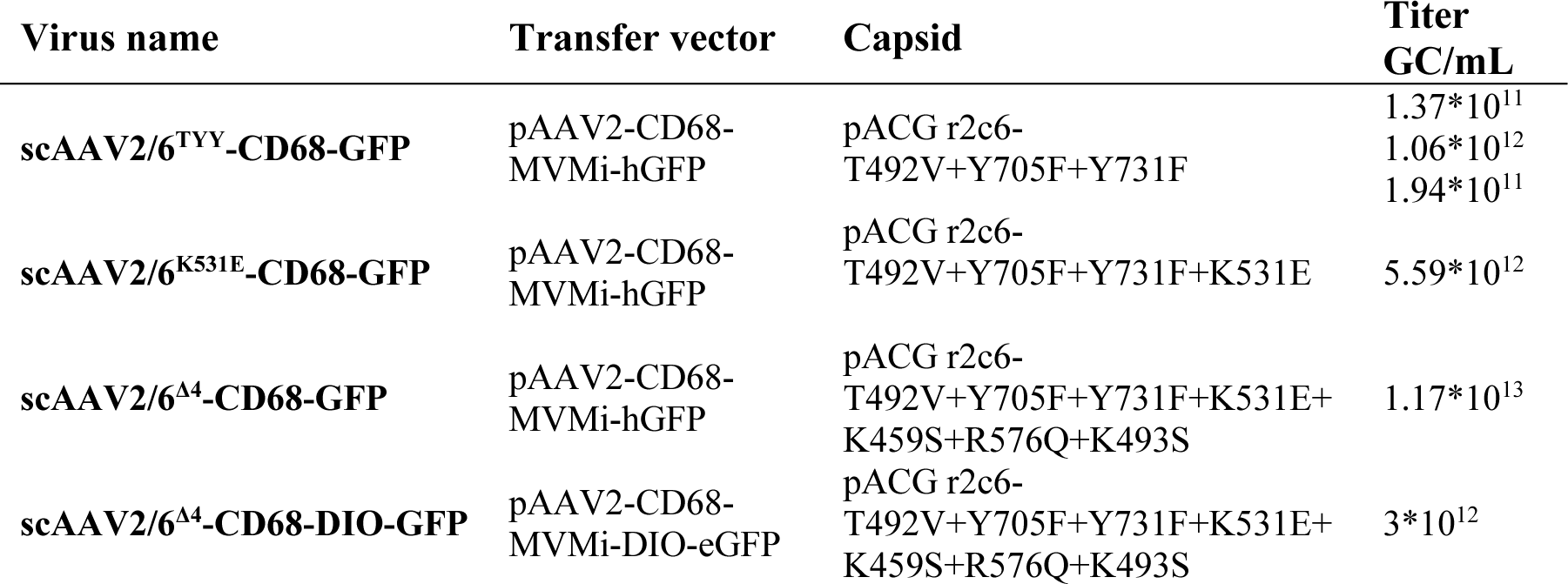
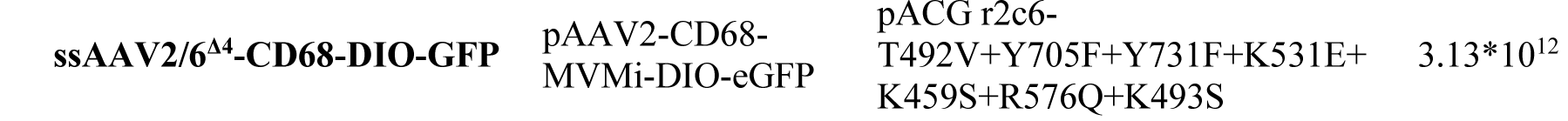
List of AAVs. Each AAV was produced with the corresponding transfer vector and capsid. Resulting titer(s) are listed (GC/mL). AAVs were diluted to titers listed for in-figure comparisons. CD68 -Cluster of Differentiation 68, CBA - Cytomegalovirus promoter and chicken beta actin enhancer, hGFP - humanized green fluorescence protein, DIO-eGFP - Double-floxed inverse orientation enhanced green fluorescence protein.

### Site directed mutagenesis

Single nucleotide substitutions were performed using the Q5® Site-Directed Mutagenesis Kit from New England BioLabs (E0552S). Exponential amplification of the viral plasmid capsid

(pACGr2c6-T492V-Y705F-Y731F) provided by *Rosario et al.* was performed according to the manufacturer’s instruction. Each mutagenesis PCR reaction was performed subsequently on the resulting plasmid from the previous confirmed mutagenesis reaction using the primer pairs listed in Table 1. Mutant vectors were transformed in chemically-competent DH5α cells (Invitrogen), and DNA isolated using Monarch mini-prep DNA kits. Mutations were confirmed by sequencing, and alignment was performed with SnapGene 4.1.9 ®.

### AAV Production

#### Transfection of HEK293T cells

HEK293T cells were purchased from the American Type Culture Collection (ATCC) and maintained at 37°C in 5% (v/v) CO2 in complete medium (DMEM high glucose GlutaMAX supplement pyruvate (Thermo Fisher Scientific, 31966047), 10% (v/v) FBS (Gibco™ Fetal Bovine Serum, qualified, E.U.-approved, South America origin, Thermo Fisher Scientific 10270106), 1% (v/v) Penicillin/Streptomycin (stock: 10,000 U/mL, Thermo Fisher Scientific, 15140122), 1% (v/v) Non-essential amino acids (Stock: 100X, Sigma-Aldrich, M7145-100ML)). Ten T150 flasks were seeded to reach 80% confluency for the day of transfection. High yield of plasmid DNA was obtained using the NucleoBond® Xtra Maxi Plus EF (Macherey-Nagel, 740426.50). 70μg AAV packaging plasmid, 70μg AAV vector plasmid and 200μg Helper plasmid were added to 50mL DMEM without serum, followed by 1360µL PEI (Polyethylenimine, Polysciences, 24765-2). 5 mL of DNA-transfection mixture was added to each T150 flask after a 15 minute incubation.

#### AAV isolation

60 hours after transfection the cells were dislodged, pelleted and stored at -80°C. For AAV isolation, cells were resuspended in lysis buffer (150mM NaCl, 20mM Tris pH 8.0, sterile filtered) and subject to three rounds of freeze/thaw cycles between dry ice/ethanol bath and 37°C water bath. MgCl2 was added (final concentration 1mM), followed by Turbonuclease (final concentration 250U/mL, BPS Bioscience, BPS 50310) to remove contaminating plasmid and genomic DNA. The cell suspension was spun down at 4,000 rpm at 4°C for 20 minutes, at which point the viral fraction was in the supernatant.

The virus was purified by discontinuous iodixanol gradient ultracentrifugation ^62^. Optiseal tubes (Beckman Coulter, 361625) were filled with a density gradient of 60%, 40%, 25% and 17% iodixanol solutions (Optiprep Iodixanol, Progen Biotechnik, 1114542).

The viral lysate supernatant was loaded on the top layer and the tubes were centrifuged at 242,000 xg at 16°C for 90 minutes in a Beckman Optima XPN-100 ultracentrifuge, using a 70Ti rotor. The AAV particles were harvested from the intersection of 60% and 40% gradients, and purified and concentrated using Amicon filters (Millipore Amicon 100K, Merck, UFC910008). 20µL aliquots were stored at -80°C and a 5µL aliquot was reserved for titration by qPCR. The DNAse (New England BioLabs, M0303S) treated virus aliquot was serially diluted (1:10 – 1:100,000) and run alongside a linearized standard template DNA (1x10^10^ – 1x10^3^ genome copies). The Luna universal qPCR Master mix (New England BioLabs, M3003L) was prepared according to the manufacturer’s instructions with a reaction volume of 10µL. and run on a BioRad C1000 cycler using the following primers: Forward – 5’CCAGCCATCTGTTGTTTGC3’, Reverse – 5’ACAATGCGATGCAATTTCC3’. The viral genome copy number per milliliter (GC/mL) was calculated as previously described by Pfaffl ^63^.

### Primary mixed glia culture

Mixed glia culture were prepared as detailed by *Bronstein et al*. ^64^. Briefly, cortices were dissected from 3-5 murine pups aged P0-P2 in ice-cold Hank’s buffered saline, then digested in 0.05% Trypsin + EDTA (1x) for 15 minutes at 37°C. The digestion was neutralized by adding (v/v) serum-containing medium (DMEM, 10% FBS, 1% Penicillin/Streptomycin, 1% Non-essential amino acids), and the cells were pelleted at 500xg for 5 mins. After one wash, the cell pellet was resuspended in 15 mL of medium, passed through a 40µm cell strainer.

This cell suspension was plated directly onto an ibidi 8-well chamber slide (200 µl/well). The culture medium was replaced after the third day and at day 10 the mixed glia culture was mature for further experiments. For viral transduction, 1e8 viral genome copies were added per well of an 8-well ibidi chamber slide (growth area 1cm^2^).

### Animals

As indicated throughout the study, mice of both sexes and ages (4-17 weeks) were used. Founder animals were purchased from the Jackson Laboratories for the following strains: C57BL/6J (#000664), Pde6b^rd10/rd10^ (#004297), Cx3cr1^creERT2^ (#020940) Cx3cr1^GFP^ (#005582)^35^ Pde6b^rd10/rd10^, Cx3cr1^GFP^ and Cx3cr1^creERT2^ mice were backcrossed to the C57BL6/J background for at least 10 generations. Animals were housed and maintained in the IST Austria Preclinical Facility, with 12 hour light-dark cycle, food and water provided *ad libitum*. All animal procedures are approved by the “Bundesministerium für Wissenschaft, Forschung und Wirtschaft (bmwfw) Tierversuchsgesetz 2012, BGBI. I Nr. 114/2012 (TVG 2012) under the number GZ BMWFW-66.018/005-WF/V3b/2016. For tamoxifen administration, Cx3cr1^creERT2/+^ and C57BL/6J mice received intraperitoneal (IP) injections of Tamoxifen (Sigma Aldrich, T5648-5G) dissolved in corn oil (Sigma Aldrich, C8267-500ML, 150mg/kg body weight, 20mg/mL stock solution) at the age of 4-6 weeks once per day for three consecutive days.

### Anesthesia and surgical preparation

Mice were anesthetized with 5% (v/v) isoflurane (Zoetis) supplemented with oxygen at a flow rate of 0.6L/min. The anesthetized mice were transferred to a heating pad placed under a Leica dissection microscope housed in a biosafety cabinet and subsequently maintained at 2.5% (v/v) isoflurane supplemented with oxygen via a nose cone during the procedure.

Proparacaine (0.5% HCl) eye drops (Ursapharm Arzneimittel GmbH) were applied to numb the eyes, and subcutaneous injection of 100µL Metacam (Meloxacam, Boehringer Ingelheim) per 25g mouse (5mg/kg) alleviated pain.

### Ocular injections

A jeweler’s forceps was used to grasp the conjunctiva, then the sclera was carefully punctured with the bevel of a 27G (Henry Schein Medical) needle just below the limbus. A Nanofil syringe equipped with a 35G blunt ended needle (World Precision Instruments) was inserted via the pre-punctured hole. For subretinal injections, the syringe needle was inserted with care to avoid the lens, and continued until resistance could be detected indicating passage through the retinal tissue. A slight retraction of the needle allows the syringe content to be released into the subretinal space of the inferotemporal or superotemporal quadrant for the right or left eyes respectively. For intravitreal injections, the needle was inserted 1-2mm into the eye. Once inserted in either method, 1µL virus was slowly released over 30 seconds into the subretinal space or into the vitreous body and the syringe remained in position for an additional 45 seconds. Triple antibiotic ointment was applied to eye after the procedure.

### Optic nerve crush

The lateral canthus of the left eye was pinched for 10 seconds using a hemostat, then a lateral canthotomy was performed to allow visualization of the posterior pole. A jeweler’s forceps was used to firmly securely the eye at the limbus of the conjunctiva. A micro-dissection scissors was used to cut the conjunctiva in both the superior and inferior direction. To expose the optic nerve, a window was created by carefully dissecting the surrounding muscle and fascia. The optic nerve was then pinched 1mm from the posterior pole for 4 seconds using a curved N5 self-closing forceps (Dumont). Triple antibiotic ointment was applied to the eye to prevent infection.

### Retina preparation and immunostaining

Following cervical dislocation and decapitation, eyes were enucleated with curved forceps. Retinas were rapidly dissected in 1X phosphate-buffered saline (PBS) and transferred to 4% (w/v) paraformaldehyde (Sigma Aldrich, P6148-1KG) for 30 mins fixation. After 3x wash in 1X PBS, retinas were placed overnight at 4°C in 30% (w/v) Sucrose (Sigma Aldrich, 84097-1KG)/PBS. After three freeze-thaw cycles on dry ice, retinas were washed three times with 1X PBS, and blocked for 1 hour at room temperature in blocking solution (1% (w/v) bovine serum albumin (Sigma A9418), 5% (v/v) Triton X-100 (Sigma T8787), 0.5% (w/v) sodium azide (VWR 786-299), and 10% (v/v) serum (either goat, Millipore S26, or donkey, Millipore S30).

For immunostaining, primary antibodies were diluted in antibody solution containing 1% (w/v) bovine serum albumin, 5% (v/v) triton X-100, 0.5% (v/v) sodium azide, 3% (v/v) goat or donkey serum for at least three days at 4°C on a shaker. The dilution factors of the antibodies are shown in Table 3. After washing, the retinas were incubated light-protected with secondary antibodies (Table 3) diluted in antibody solution for 2 hours at room temperature on a shaker. The retinas were washed three times with 1X PBS for 30 minutes. The nuclei were labelled with Hoechst 33342 (1:5,000, Thermo Fisher Scientific, H3570) in 1X PBS for 10 minutes at RT and washed again three times with 1X PBS for 30 minutes. The retina was whole mounted on a glass cover slide with the ganglion cell layer facing up, Antifade solution containing 10% (v/v) mowiol (Sigma, 81381), 26% (v/v) glycerol (Sigma, G7757), 0.2M tris buffer pH 8, 2.5% (w/v) Dabco (Sigma, D27802) was added and the retinas were coverslipped (#1.5 VWR, 631-0147).

**Table 3:**
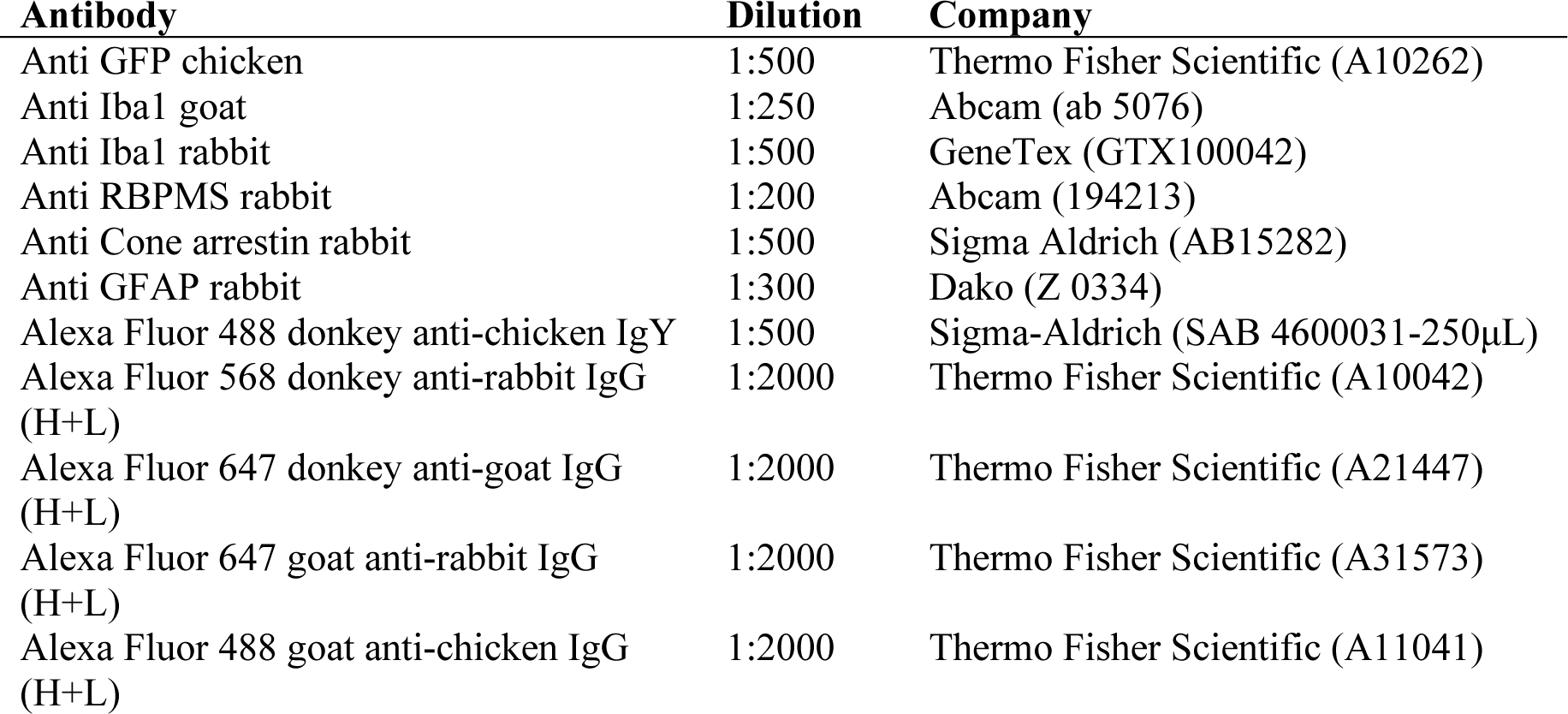
Dilution table of primary and secondary antibody including vendor, and catalog number.

### Confocal microscopy and image analysis

Flat mounted retinas were imaged with an Axio Imager Z2 Zeiss LSM800 upright confocal microscope using a Nikon Plan-Apochromat 20X magnification air objective (NA 0.8). A 2x2 tile scan image was acquired in two ROIs of retina measuring 0.312 x 0.312 μm with Nyquist z-steps. All images were acquired using the same settings. Stitched tile scans were analyzed in Imaris v9.3 using the spots function to facilitate cell counting. Transduction efficiency of microglia was analyzed in the outer and the inner plexiform layers of the retina. Transduction efficiency was calculated by dividing the number of transduced Iba1^+^ cells by the total number Iba1^+^ cells within a scanned ROI. Sholl analysis was determined by the number of filament sholl intersections exported from a 3-dimensional microglial trace using the Filament tracing plug-in in Imaris v9.3.

### Statistical analysis

All statistics were performed using the statistical functions in SciPy library (v1.6.2) in python as indicated in the figure legends. Retinas were excluded from analysis if a cataract was present at the time of retinal dissection or if high macrophage infiltration in the tissue was observed, indicating significant tissue damage from the injection. Error bars represent the standard error of the mean.

## Author contributions

MM and SS conceived and developed experimental design and wrote the manuscript. MM, GW, GC and RC performed experiments. MM and GW analyzed data, performed statistics and created figures. All authors and read and approved the final manuscript.

## Acknowledgements

This project has received funding from the European Research Council (ERC) under the European Union’s Horizon 2020 research and innovation programme (grant agreement No 715571). The research was supported by the Scientific Service Units (SSU) of IST Austria through resources provided by the Bioimaging Facility, the Life Science Facility, and the Pre-Clinical Facility, namely Sonja Haslinger and Michael Schunn for their animal colony management and support. We would also like to thank Chakrabarty Lab for sharing the plasmids for AAV2/6 production. Finally, we would like to thank the Siegert team members for the discussion about the manuscript.

## Supplementary Figures

**Supplementary Figure 1.**
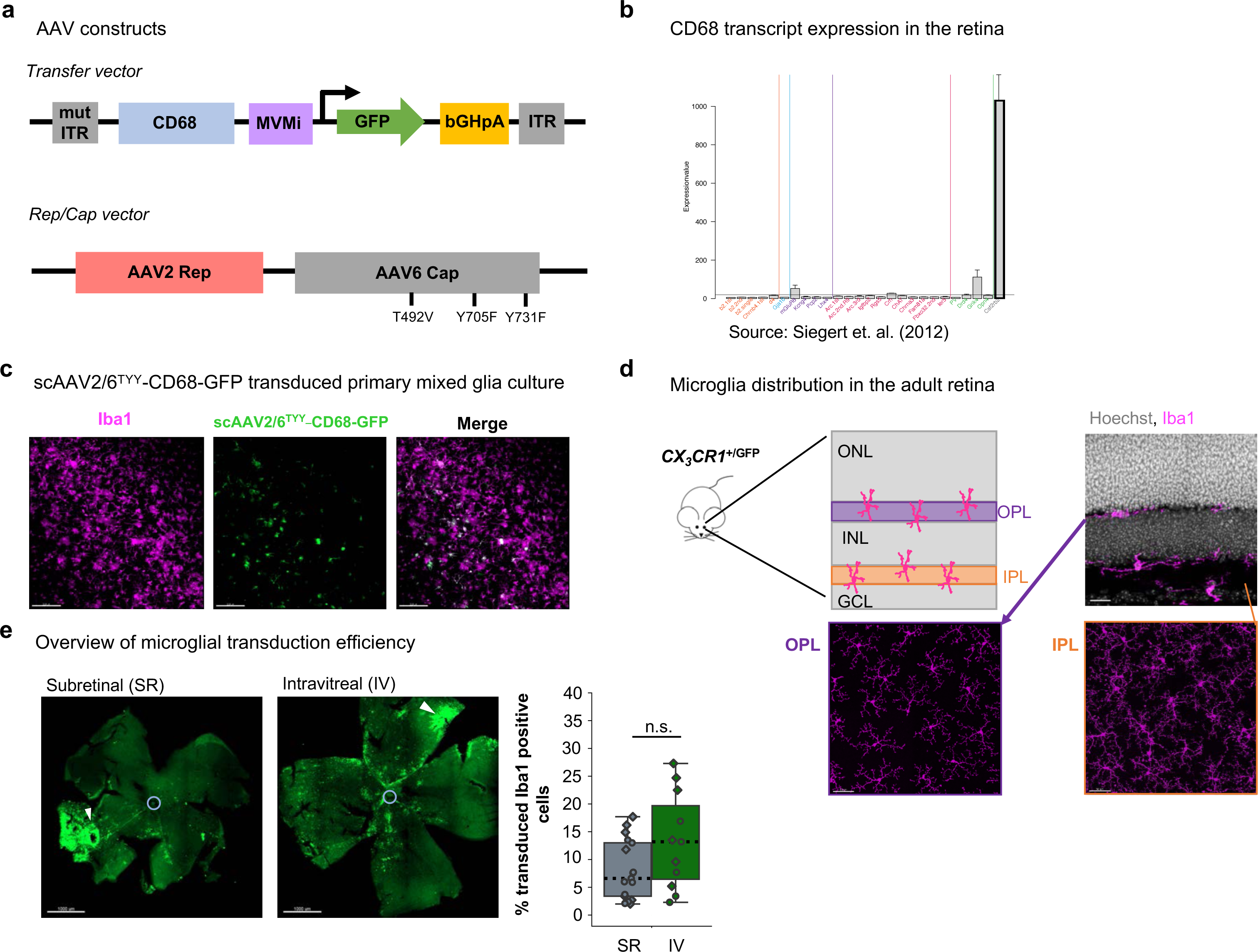
scAAV2/6^TYY^-CD68-GFP design and validation *in vitro* and *in vivo*. (a) Transfer vector illustration containing the CD68 promoter, MVM intron and Rep/Cap vector with indicated TYY mutations in capsid sequence. (b) CD68 transcript abundance in retinal cell types of the retina. https://www.fmi.ch/roska.data/index.php. ^6^. Outlined bar: microglia. (c) Mixed primary glia cultures transduced with 1*10^8^ viral genomes/well of scAAV2/6^TYY^-CD68-GFP (green) counterstained with Iba1 (magenta). Scale bar: 200µm. (d) Pictorial representation of the retinal layers, highlighting the outer plexiform layer (OPL, purple) and inner plexiform layer (IPL, orange), with corresponding confocal images of retinal cross section or whole mount images of Iba1-stained microglia (magenta). Scale bar: 50µm. (e) Overview images of retinas from subretinal (SR) or intravitreal (IV) injection indicating viral-mediated GFP transgene expression. Blue circle indicates optic nerve head. White triangles: injection site. Total microglial transduction efficiency quantification from ROI1 using 20X images. (Wilcoxon rank-sum test: *P* = 0.011). Scale bar: 1000µm. \**P* < 0.05, ^ns^*P* > 0.05. ITR, inverted terminal repeat; CD68, Cluster of differentiation 68; MVMi, minute virus of mice intron; GFP, green fluorescent protein; bGHpA, bovine growth hormone polyA; ONL, outer nuclear layer; INL, inner nuclear layer; GCL, ganglion cell layer.

**Supplementary Figure 2.**
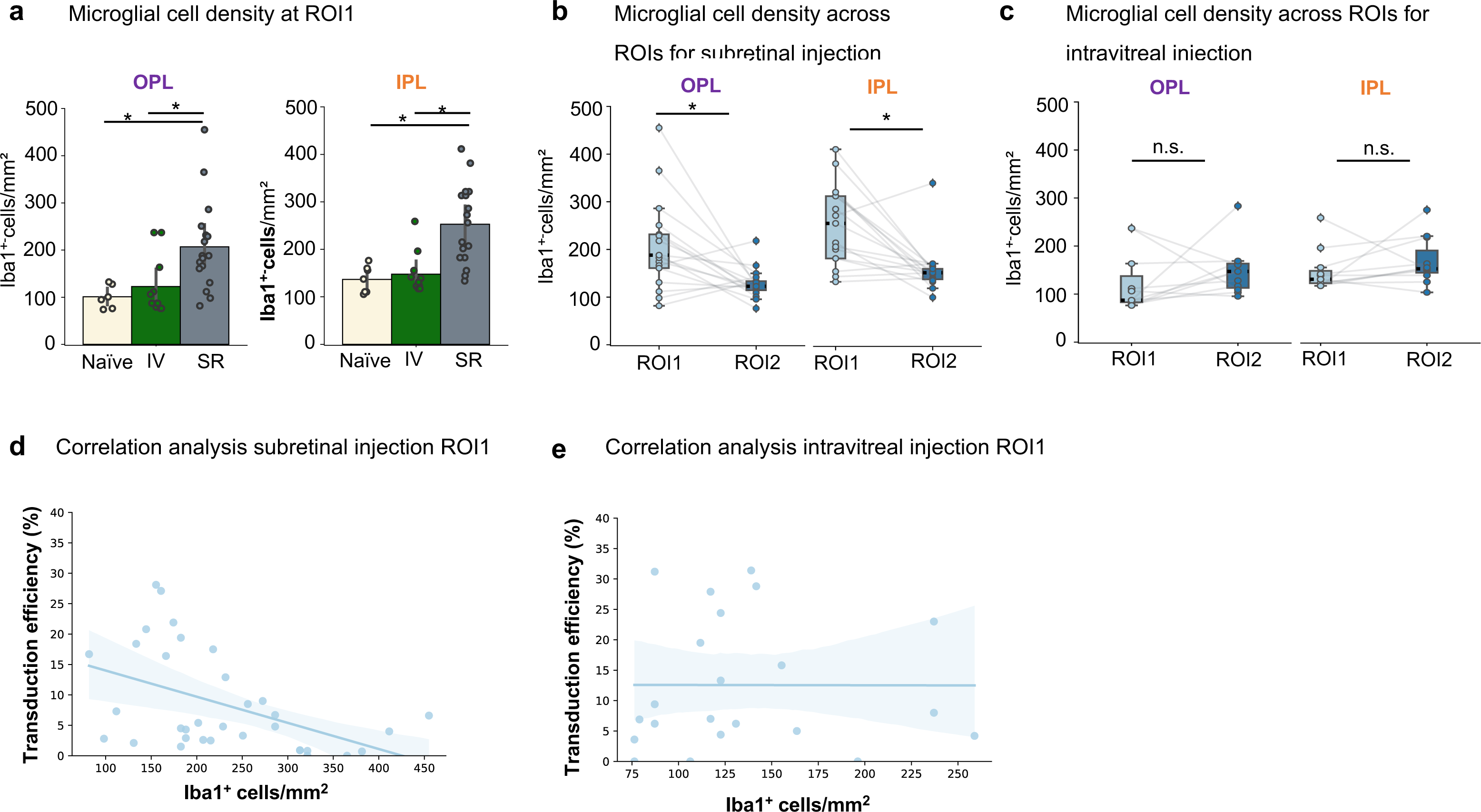
Microglial cell density after viral injection. (a) Quantification of OPL_microglia_ and IPL_microglia_ cell densities per mm^2^ in retinas from naïve (un-injected), intravitreal or subretinal injections at ROI1. (Kruskal-wallis test: OPL, *P* = 0.0002; IPL, *P* = 0.003; post-hoc comparisons Wilcoxon rank-sum test: OPL SR-Naïve, *P* = 0.002; OPL SR-IV, *P* = 0.0004; IPL SR-Naïve, *P* = 0.004; IPL SR-IV, *P* = 0.010). (b, c) Comparison of microglial density between ROIs for both OPL and IPL niche after subretinal (Wilcoxon signed-rank test: OPL*, P* = 0.004; *IPL, P* = 0.004) or intravitreal injections (c, Wilcoxon signed-rank test: OPL*, P* = 0.248; *IPL, P* = 0.153). (d, e) Correlation using linear regression model between microglial cell density and transduction efficiency for subretinal (*Pearson’s r* = -0.476, *P* = 0.004) or (e) intravitreal injection (*Pearson’s r* = 0.002, *P* = 0.992). \**P* < 0.05, ^ns^*P* > 0.05.

**Supplementary Figure 3.**
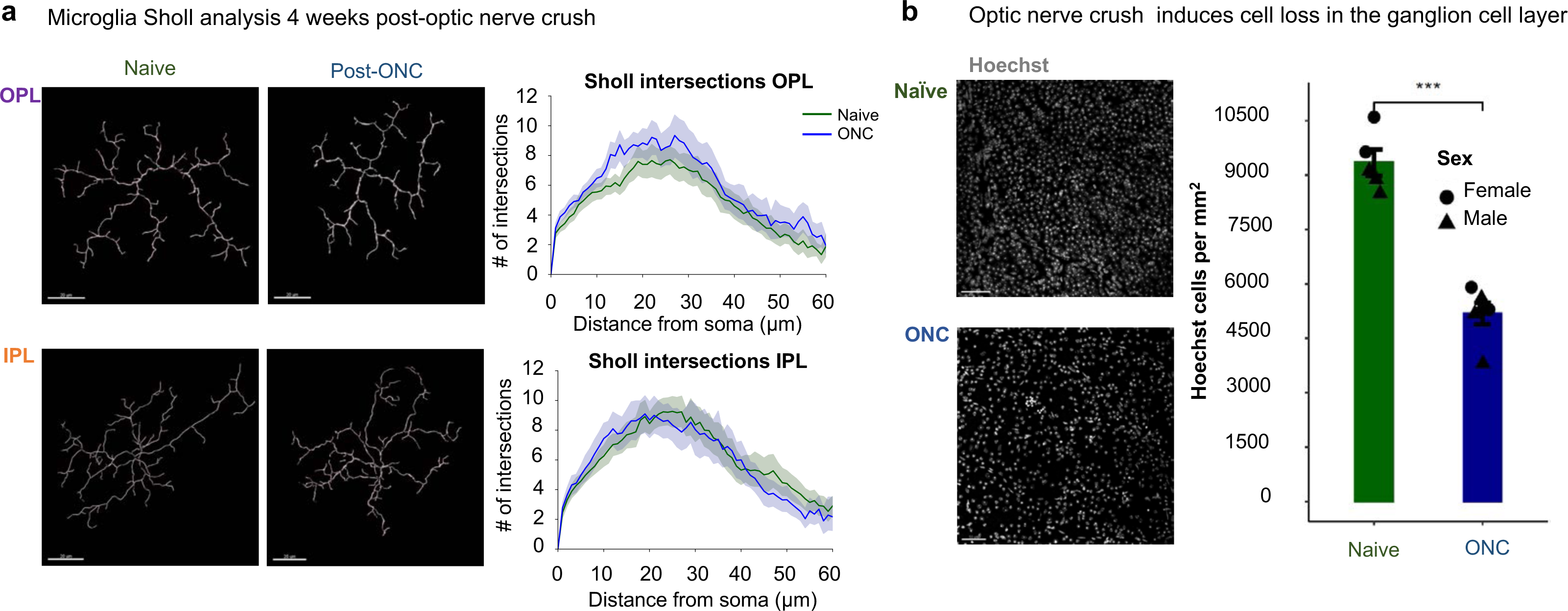
Microglial morphology and ganglion cell loss four weeks after optic nerve crush. (a) Example microglial tracings for naïve or four weeks post-optic nerve crush in OPL or IPL. Sholl intersections quantification from 40x 2x2 tiled images from >18 cells from at least 2 animals for each layer and condition. Scale bar: 20µm. (b) Hoechst-stained retinal whole mount images of ganglion cell layer (GCL). Total number of nuclei in the GCL is reduced in ONC. Scale bar: 50µm (Welch’s Two-sample t-test: *P* = 0.00001). ***P< 0.005

**Supplementary Figure 4.**
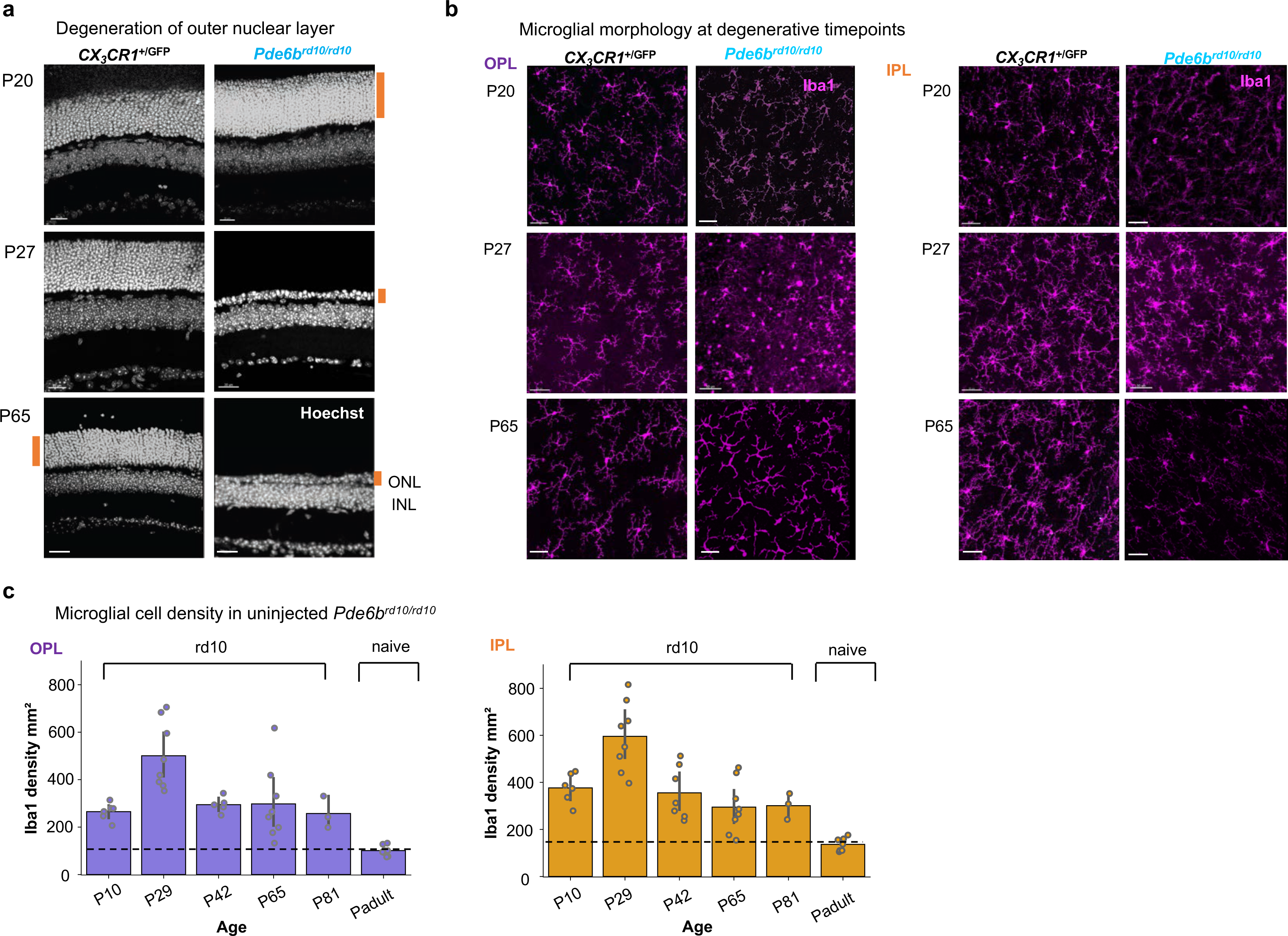
ONL thinning and microglial phenotype in *Pde6b^rd10/rd10^* degeneration model. (a) Hoechst-labeled retinal cross sections from control Cx3Cr1^GFP/+^ or *Pde6b^rd10/rd10^* mice at P20, P27 and P65. Orange bar highlights ONL thickness. Scale bar: 20µm. (b) Retinal wholemounts immunostained with Iba1 (magenta) depicting OPL_microglia_ or IPL_microglia_ in control Cx3Cr1^GFP/+^ or *Pde6b^rd10/rd10^* mice. Scale bar: 50µm. (c) Microglial cell density in OPL or IPL of uninjected *Pde6b^rd10/rd10^* compared to naïve (dashed line) adult mice. (d) Cell density at ROI1 of OPL or IPL microglia in P65 *Pde6b^rd10/rd10^* retinas injected with scAAV2/6^TYY^-CD68-GFP. ONL, outer nuclear layer; INL, inner nuclear layer.

**Supplementary Figure 5.**
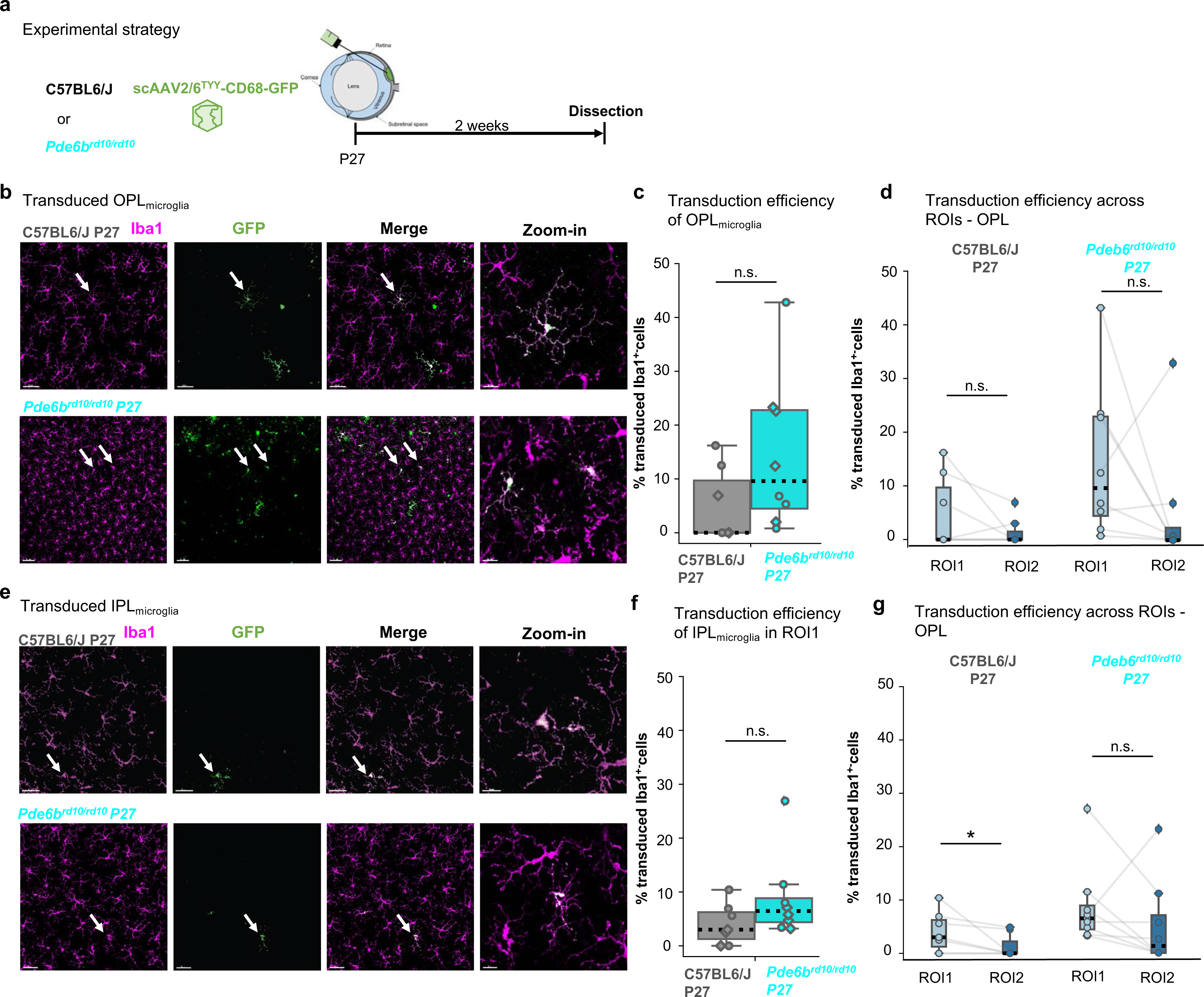
ONL loss in P27 *Pde6b^rd10/rd10^* benefits OPL microglia. (a) Experimental timeline. *Pde6b*^rd10/rd10^ or C57BL6/J mice received subretinal injection of scAAV2/6^TYY^-CD68-GFP (1.37*10^11^gc/mL) at postnatal day 27 (P27) and retinas were collected 2 weeks after injection. (b, e) Retinal wholemounts of OPL_microglia_ (b) and IPL_microglia_ (e) from indicated strain after subretinal injection stained with Iba1 (magenta) and GFP (green). White arrows indicate zoom-in region. Scale bar: 50µm, zoom-in: 15µm. (c, f) Percent transduction efficiency in OPL_microglia_ (c, Wilcoxon ranked-sum test, *P*=0.105) and IPL_microglia_ (f, Wilcoxon ranked-sum test, *P*=0.132) Stats. (d, g) Transduction across ROIs in C57BL6/J and P27 *Pde6b^rd10/rd10^* retinas in the OPL (d, Wilcoxon signed-rank test: C57BL6/J*, P* = 0.144; *Pde6b^rd10/rd10^, P* = 0.123) and IPL (g, Wilcoxon signed-rank test: C57BL6/J*, P* = 0.043; *Pde6b^rd10/rd10^, P* = 0.161). *Pde6b*^rd10/rd10^: n= 8 retinas, 7 mice. C57BL6/J: n= 7 retinas, 4 mice. \**P* < 0.05, ^ns^*P* > 0.05.

**Supplementary Figure 6.**
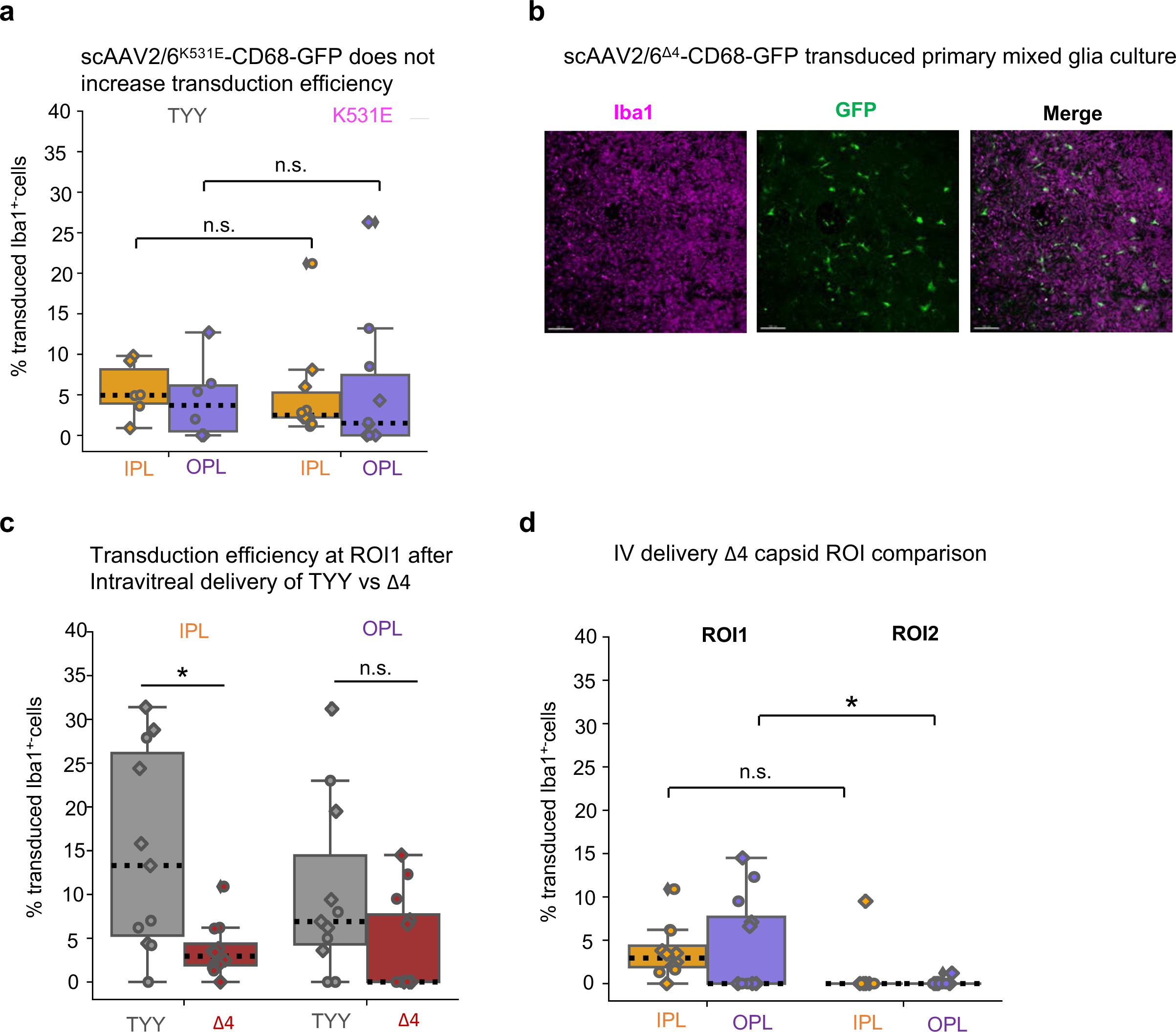
Single heparin-binding mutant and intravitreal delivery of AAV2/6 ^Δ4^. (a) Single capsid mutant scAAV2/6^K531E^-CD68-GFP microglial transduction efficiency compared to scAAV2/6^TYY^-CD68-GFP (Wilcoxon ranked-sum test, OPL, *P* = 0.828; IPL, *P =* 0.329). (b) Mixed primary glia cultures transduced with 1*10^8^ viral genomes of scAAV2/6^Δ4^-CD68-GFP counterstained with Iba1 (magenta). Scale bar: 200 µm. (c) Comparison of intravitreally delivered scAAV2/6^TYY^-CD68-GFP and scAAV2/6^Δ4^-CD68-GFP transduction efficiency of OPL_microglia_ and IPL_microglia_ (Wilcoxon ranked-sum test: OPL, *P* = 0.139; IPL, *P*=0.005). (d) Comparison across ROIs for both OPL and IPL niche after intravitreal delivery of scAAV2/6^Δ4^-CD68-GFP. TYY: 11 retinas, 6 mice. Δ4: 12 retinas, 6 mice. \**P* < 0.05, ^ns^*P* > 0.05.

**Supplementary Figure 7.**
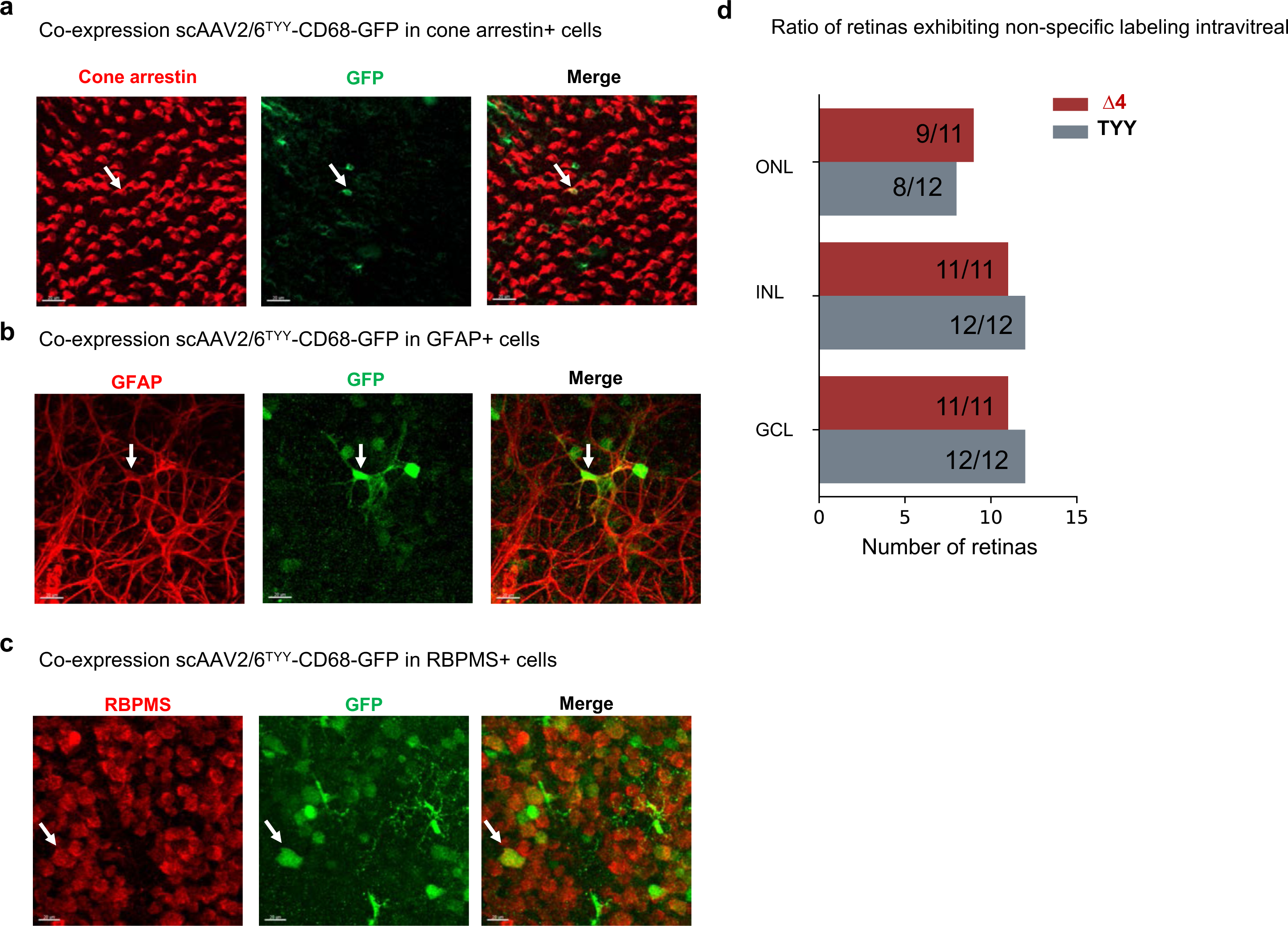
Non-specific labeling using scAAV2/6^TYY^-CD68-GFP Retinal wholemounts of. (a) cone photoreceptors (cone arrestin^+^, red), (b) astrocytes (GFAP^+^, red), (c) retinal ganglion cells (RPBMS^+^) co-expressing GFP (green) following subretinal injection of scAAV2/6^TYY^-CD68-GFP. White arrow indicates co-localized cell. Scale bar: 50µm. (d) Ratio of analyzed retinas exhibiting non-specific GFP expression after subretinal delivery of scAAV2/6^TYY^-CD68-GFP across the nuclear layer, outer nuclear layer (ONL), inner nuclear layer (INL) and ganglion cell layer (GCL).

**Supplementary Figure 8.**
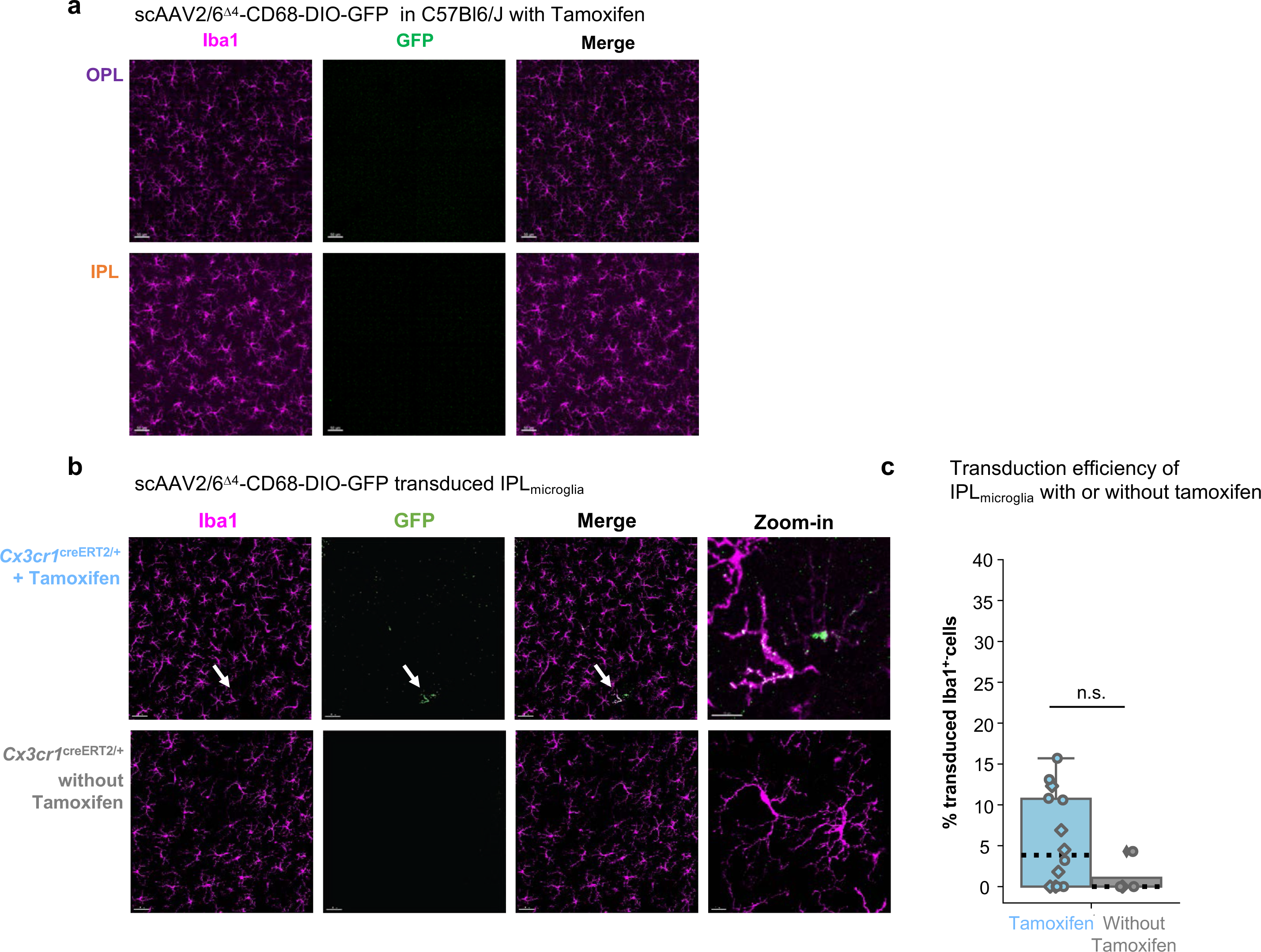
IPL_microglia_ transduction after scAAV^Δ4^-CD68-DIO-GFP delivery. (a) Retinal wholemounts of OPL and IPL microglia of C57BL6/J mice after subretinal delivery of scAAV^Δ4^-CD68-DIO-GFP and Tamoxifen treatment immunostained with Iba1 (magenta) and GFP (green). Scale bar: 50µm. (b) Retinal wholemounts of IPL_microglia_ in Cx3cr1^creERT2/+^ mice after subretinal delivery of scAAV^Δ4^-CD68-DIO-GFP with or without Tamoxifen induction immunostained with Iba1 (magenta) and GFP (green). White arrows indicate zoom-in. Scale bar: 50µm, zoom-in: 15µm. (c) Comparison of transduction efficiency in the IPL with and without Tamoxifen (Wilcoxon ranked-sum test: *P* = 0.151). C57BL6/J: 12 retinas, 7 mice. Cx3cr1^creERT2/+^ with Tam: 14 retinas, 9 mice. Cx3cr1^creERT2/+^ without Tam: 4 retinas, 3 mice. \**P* < 0.05, ^ns^*P* > 0.05.

